# Comprehensive interrogation of a *Drosophila* embryonic patterning network reveals the impact of chromatin state on tissue-specific burst kinetics and RNA Polymerase II promoter-proximal pause release

**DOI:** 10.1101/2022.10.25.513691

**Authors:** George Hunt, Roshan Vaid, Sergei Pirogov, Alexander Pfab, Christoph Ziegenhain, Rickard Sandberg, Johan Reimegård, Mattias Mannervik

**Affiliations:** Dept. Molecular Biosciences, The Wenner-Gren Institute, Stockholm University, Stockholm, Sweden; Dept. Cell and Molecular Biology, Karolinska Institutet, Sweden; Dept. Cell and Molecular Biology, National Bioinformatics Infrastructure Sweden, Science for Life Laboratory, Uppsala University, Uppsala, Sweden

## Abstract

Formation of tissue-specific transcriptional programs underlies multicellular development, but how the chromatin landscape influences transcription is not fully understood. Here we comprehensively resolve differential transcriptional and chromatin states during *Drosophila* dorsoventral (DV) patterning. We find that RNA Polymerase II pausing is established at DV promoters prior to zygotic genome activation (ZGA), that pausing persists irrespective of cell fate, but that release into productive elongation is tightly regulated and accompanied by tissue-specific P-TEFb recruitment. DV enhancers acquire distinct tissue-specific chromatin states through CBP-mediated histone acetylation that predict the transcriptional output of target genes, whereas promoter states are more tissue invariant. Transcriptome-wide inference of burst kinetics in different cell types revealed that while DV genes are generally characterized by a high burst size, either burst size or frequency can differ between tissues. The data suggest that pausing is established by pioneer transcription factors prior to ZGA and that release from pausing is imparted by enhancer chromatin state to regulate bursting in a tissue-specific manner in the early embryo. Our results uncover how developmental patterning is orchestrated by tissue-specific bursts of transcription from Pol II primed promoters in response to enhancer regulatory cues.

## Introduction

The ability to dynamically regulate gene expression is integral to developmental processes in multicellular organisms by enabling cells that retain identical DNA sequences to form specialized cell types. Early *Drosophila* embryogenesis involves 13 rapid, synchronous nuclear divisions within a syncytium to give rise to ~6000 nuclei that then cellularize, undergo zygotic genome activation (ZGA), and become specified. Dorsoventral (DV) axis specification of the early *Drosophila* embryo is one of the most well studied gene regulatory networks ^1,2^. During DV patterning, distinct cell fates form in response to an intranuclear morphogen gradient of the maternally supplied REL-family transcription factor Dorsal (Dl) ^3–5^. Differential activation of Toll receptors leads to high nuclear import of Dl in ventral regions, low levels of nuclear Dl in lateral regions and an absence of Dl in dorsal regions ^6^. The Dl gradient forms during nuclear cycles 10-14 and induces distinct complements of zygotic genes in ventral, lateral and dorsal regions of the embryo, leading to cell specification at nuclear cycle 14 and formation of presumptive mesoderm, neurogenic ectoderm and dorsal ectoderm, respectively (Fig. 1A). Dl activates genes such as *twist* (*twi*) in the mesoderm and *intermediate neuroblasts defective* (*ind*) in the neuroectoderm, but can also function as a repressor, which restricts genes such as *decapentaplegic* (*dpp*) to the dorsal ectoderm where Dl is absent from the nuclei (Fig. 1B).

**Figure 1.**
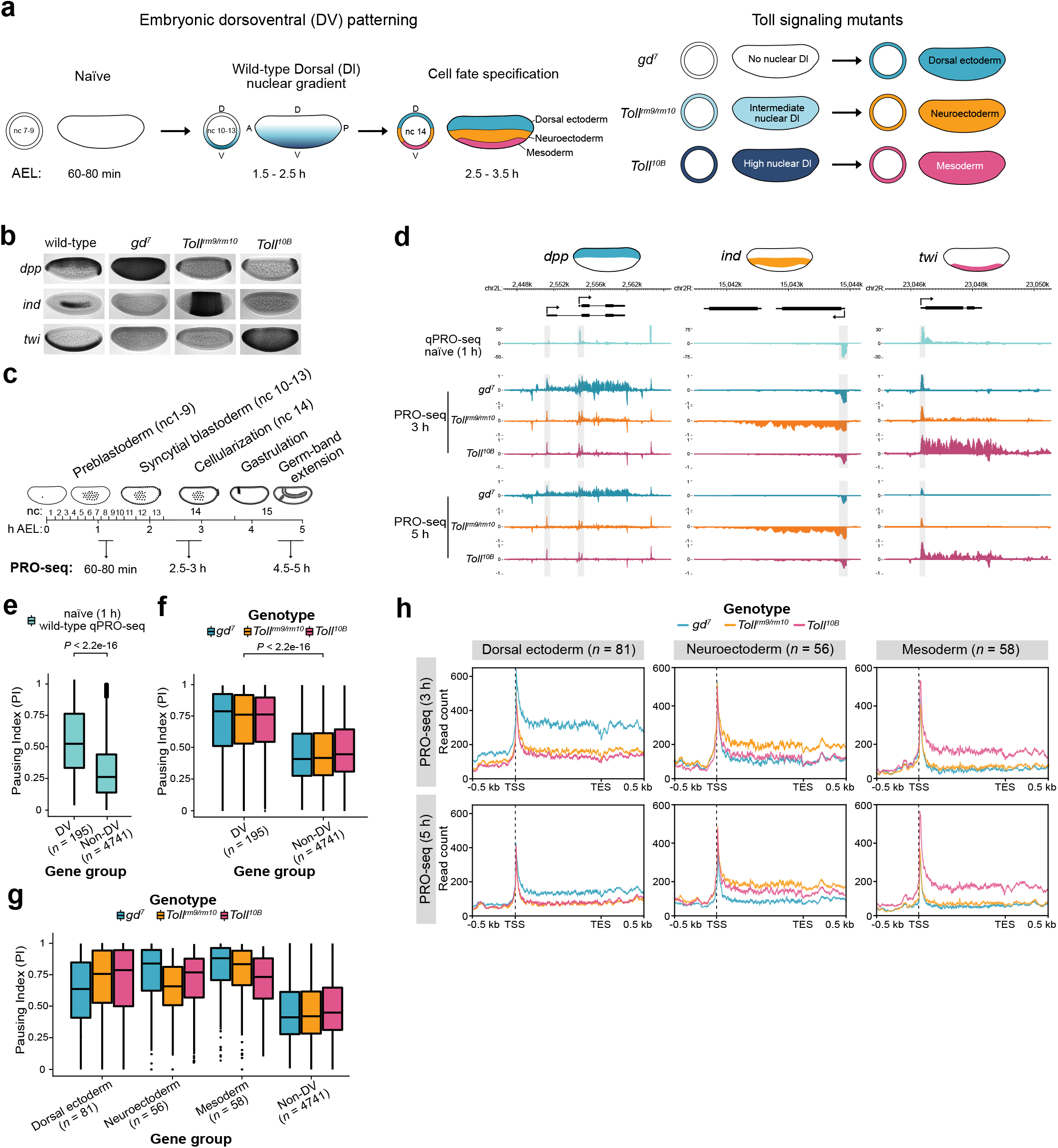
Promoter-proximal paused Pol II is established at DV regulated genes prior to ZGA but is released into elongation in a tissue-specific manner. **a)** Schematic of embryonic DV patterning. From an initially transcriptionally inert naïve embryo (nuclear cycle (nc) 7-9, 1-1.2 hours (h) after egg laying (AEL)), a dorsoventral (DV) nuclear gradient of the maternally supplied transcription factor dorsal (Dl) (nc 10-13, 1.5-2.5 h AEL) directs cell fate specifications at zygotic genome activation (ZGA) (nc 14, 2.5-3.5 h AEL). Distinct transcriptional programs initiated by the absence of Dl dorsally, moderate nuclear Dl laterally and high nuclear Dl ventrally lead to cell specification into dorsal ectoderm, neuroectoderm and mesoderm, respectively. Disrupted Dl gradient formation in *Toll* signaling mutants produces embryos composed entirely of presumptive dorsal ectoderm (*gd^7^*), neuroectoderm (*Toll^rm9/rm10^*) and mesoderm (*Toll^10B^*). **b)** Images of whole mount *in situ* hybridization in wild-type and *Toll* mutant embryos (2-4 h AEL) with probes hybridizing to mRNAs of representative DV regulated genes (*dpp*, *ind* and *twi*). **c)** Schematic of the experimental design to study spatio-temporal transcriptional dynamics during DV patterning. PRO-seq was performed on naïve wild-type embryos (nc 7-9, 60-80 min AEL) and *Toll* mutant embryos at ZGA (nc 14, 2.5-3 h AEL) and after gastrulation (> nc 14, 4.5-5 h AEL). **d)** Genome browser shots of stranded PRO-seq signal (RPKM x10^3^) at *dpp*, *ind* and *twi*. Promoters are shaded gray. **e)** Pausing index (PI) of DV and non-DV regulated genes from qPRO-seq in wild-type naïve (1 h) embryos and **(f)** PRO-seq in *Toll* mutants. **g)** PI of DV regulated genes partitioned by the tissue of expression from PRO-seq in *Toll* mutants. **h)** Metagene plots of *Toll* mutant PRO-seq (2.5-3 and 4.5-5 h) at DV regulated genes. Comparisons of the PI between DV and non-DV gene classes are from the Wilcoxon Rank-sum test.

An important aspect of transcriptional regulation is how regulatory signals are conveyed from enhancers to elicit a transcriptional response at the promoter. Hi-C, Micro-C and microscopy-based data revealed that there are no differences in the topologically associated domain (TAD) structure or enhancer-promoter (E-P) contact frequencies for DV genes between cells in the embryo where they are expressed or silent ^7,8^. This suggests that E-P looping is not the step that triggers tissue-specific activation of DV genes. Pausing of transcriptionally engaged RNA Polymerase II (Pol II) 30-60 bp downstream of the transcription start site (TSS) has been identified as an important regulatory checkpoint that allows the release of Pol II in to productive elongation to be tightly controlled ^9,10^. Pol II pausing is prevalent among developmental genes during *Drosophila* embryogenesis ^11^, and allows cells in a tissue to synchronously activate gene expression ^12^.

DV tissue mutant embryos, derived from maternal effect mutations, with either the absence (*gd^7^*, dorsal ectoderm), or uniformly low (*Toll^rm9/rm10^*, neurogenic ectoderm) and high (*Toll^10B^*, mesoderm) levels of nuclear Dl (Fig. 1a,b), provided an amenable substrate for ChIP-based approaches to characterise DV enhancers and other important regulatory elements based on the enrichment of histone modifications such as H3K27ac and occupancy of the co-activator CBP ^7,13–17^. Nonetheless, a comprehensive genome-wide assessment of the interplay between transcriptional activity and chromatin state across the DV axis is lacking.

In this study, we used the DV patterning model to examine the spatio-temporal interplay between transcription and chromatin state. We performed Precision Run-On Sequencing (PRO-seq) on precisely aged tissue mutant *Drosophila* embryos to measure nascent transcription and Pol II pausing genome-wide, alongside chromatin state data from ATAC-seq, ChIP-seq and CUT&Tag. We further inferred transcriptional burst kinetics from single-cell RNA-seq data. Our findings suggest that enhancers and promoters are initially primed for activation competency across cells that adopt distinct fates, but the spatio-temporally regulated acquisition of distinct patterns of enhancer CBP occupancy and histone acetylation in response to the Dl gradient leads to differential DV gene expression by controlling burst kinetics and the release of paused Pol II into productive elongation.

## Results

### Paused Pol II is established at dorsoventral genes prior to their expression in the early embryo

To obtain a precise genome-wide assessment of the activity state of Pol II and spatio-temporal differences in zygotic transcription during DV patterning, we performed PRO-seq on naïve wild-type embryos, 60-80 min after egg laying (AEL), and on DV tissue mutant embryos composed entirely of presumptive dorsal ectoderm (*gd^7^*), neurogenic ectoderm (*Toll^rm9/rm10^*) or mesoderm (*Toll^10B^*) at 3 and 5 hours AEL (Fig. 1a-d) ^18^. For the naïve stage, we also hand-sorted embryos to ensure that they were not older than nuclear cycle (nc) 9, and used the more sensitive qPRO-seq protocol ^19^. We identified differentially expressed genes between the mutant embryos by comparing the number of PRO-seq reads mapping to the gene body (defined as the coding sequence (CDS) of the gene), and observed 195 genes that were up-regulated specifically in one of the mutants (Fig. S1a,b and Table S1). A comparison with previously published DV regulated genes ^20^ showed a large overlap and expression in the expected tissue (Fig. S1c,d). Gene ontologies for the differentially expressed genes were consistent with their expected functions in epithelial, nervous system and muscle development, respectively (Table S2). Most DV regulated genes were expressed at both 3 and 5 h AEL, but some were specific to the later time point (Fig. S1e, Table S1).

Many developmental genes exhibit promoter-proximal paused RNA polymerase II (Pol II) ~30-60 bp downstream of the TSS ^21^. To measure pausing, we calculated the pausing index from the ratio of PRO-seq reads mapping to the promoter (from 50 bp upstream of the TSS to 100 bp downstream of the TSS) and the sum of reads mapping to the promoter and the gene body, which revealed that DV genes, as well as anterior-posterior (AP) patterning genes, were more highly paused than non-DV genes expressed in these embryos (Fig. 1e,f, S1f). Interestingly, Pol II pausing was observed at DV genes already in the naïve stage, prior to their expression (Fig. 1d,e).

To ensure that detection of paused Pol II in the naïve stage was not due to sample contamination with older embryos, we measured the gene body read counts and pausing index of zygotic genes expressed at specific stages of development ^22^ (Fig. S1g-j). Genes already expressed at nc 7-9 and nc 9-10 had higher gene body qPRO- and PRO-seq signal than DV genes and genes expressed at the syncytial (nc 11-13) and cellularized (nc 14) blastoderm stages, demonstrating that the experiments captured properly staged embryos (Fig. S1g,h). Whereas DV genes were paused at the naïve stage, genes expressed at the naïve stage had a low pausing index, consistent with previous findings ^23^ (Fig. S1j). Core promoter motifs have been shown to strongly influence Pol II recruitment and pausing ^9,24,25^. Examination of the CORE database ^26^ and *de novo* motif analysis showed that DV genes were highly enriched for core promoter motifs ^27^, such as Initiator (Inr), downstream promoter element (DPE) and TATA-box, compared to other genes (Fig. S1k-n and Table S3), likely contributing to their high pausing index.

High Pol II pausing was maintained at dorsal ectoderm, neuroectoderm, and mesoderm-specific genes across all three DV mutants (Fig. 1f), but gene body reads were elevated in specific mutants, as exemplified by *decapentaplegic* (*dpp), intermediate neuroblasts defective (ind)* and *twist* (*twi)* (Fig. 1d). Similar results were obtained with Pol II antibodies in CUT&Tag on *Toll* mutant embryos (Fig. S1o). The pausing index for DV genes was lower in the tissue mutant of expression (Fig. 1g). To address whether the reduction in pausing was due to the elevated gene body reads in the tissue of expression, or a decrease in reads for promoter-proximal paused Pol II, we measured the signal for these regions separately for all genes (Fig. S1p, Table S1), and generated metaplots of PRO-seq read density (Fig. 1h). The promoter-proximal Pol II signal was similar among the three mutants for most genes at 3 h (AEL), and the reduced pausing index was mostly explained by the elevation of gene body reads, suggesting a key role for pause release in DV gene transcription. The observation that DV genes become highly paused in naïve embryos prior to their transcription and that pausing is maintained in different tissue contexts, irrespective of transcription, demonstrates that pause release is a major regulatory step in tissue-specific DV transcription.

### Enhancer chromatin state reflects tissue-specific DV gene transcription

To identify what controls the release of paused Pol II into productive elongation, we examined the chromatin states of enhancers and promoters for DV genes. Occupancy of p300/CBP and enrichment of the p300/CBP catalyzed mark H3K27ac are hallmarks of active enhancers ^28–31^ and DV enhancers have previously been identified based on differential H3K27ac ^7,14^. We screened for DV enhancers by correlating differential expression with genomic regions that exhibit tissue-specific *Drosophila* CBP (Nejire) occupancy, H3K27ac enrichment and chromatin accessibility (ATAC-seq) (Fig. 2a).

**Figure 2.**
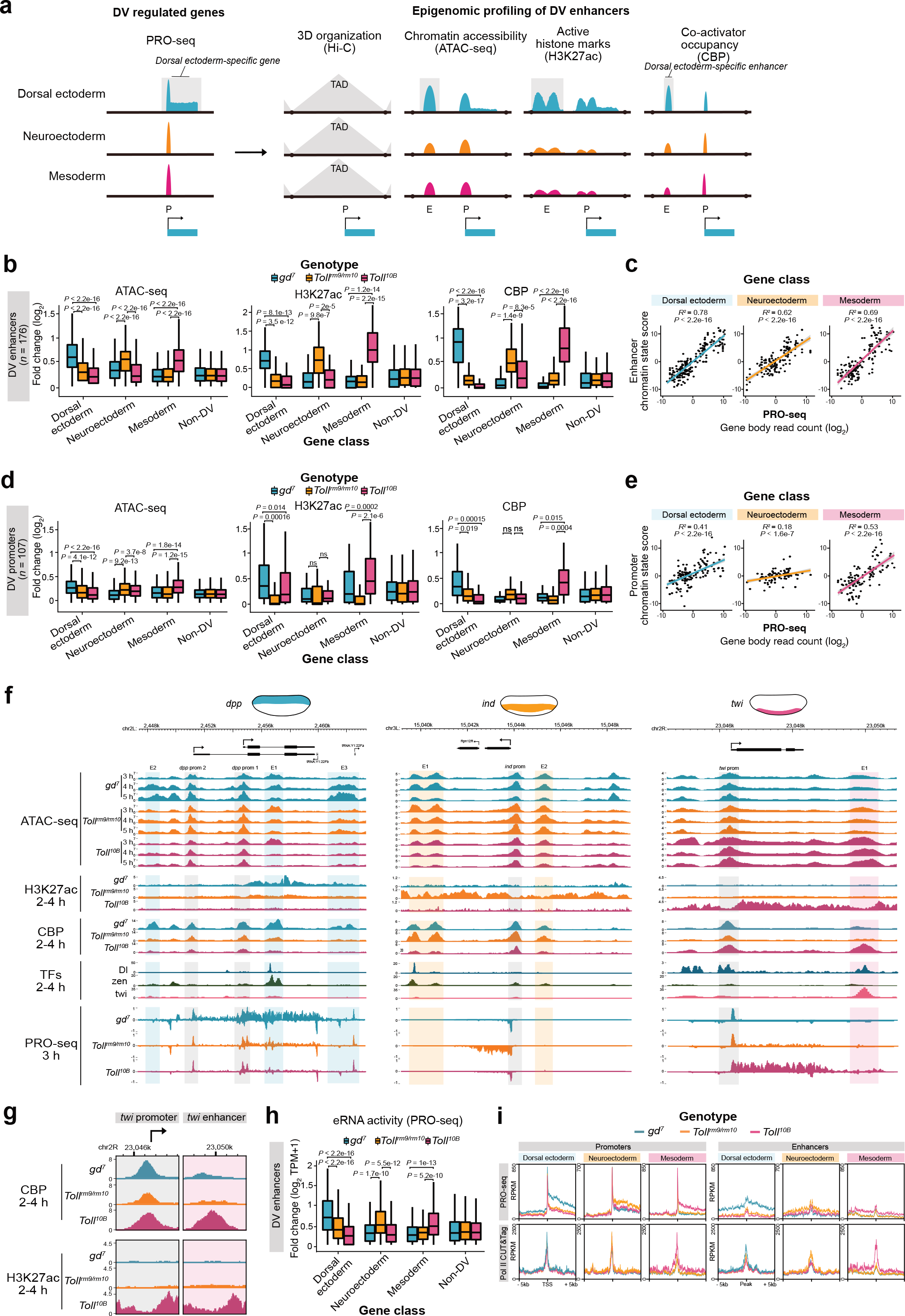
Epigenomic profiling identifies chromatin states at DV enhancers and promoters that correlate with tissue-specific gene expression. **a)** Schematic of the epigenomic profiling strategy for identifying tissue-specific DV enhancers genome-wide. PRO-seq identified DV genes were linked to regions within the same topologically associating domain (TAD) with differential chromatin accessibility (ATAC-seq), enrichment of the active histone mark H3K27ac and occupancy of CBP between *Toll* mutants. **b)** Boxplots of the fold change (log_2_) ATAC-seq, CBP and H3K27ac enrichment between *Toll* mutants at DV enhancers separated by the tissue of expression of their target genes. **c)** The correlation between the combined tissue-specific enhancer chromatin state score and target DV gene expression (PRO-seq gene body count (GBC)) for each tissue-specific gene classes. The *P*-value from Pearson’s correlation is shown alongside the coefficient of determination (*R^2^*). **d)** The fold change (log_2_) in enrichment between *Toll* mutants for the genomic datasets in **b** at promoters associated with DV enhancers. **e)** Correlations between the combined tissue-specific promoter chromatin state score and gene expression (PRO-seq, GBC). **f)** Genome browser shots of *Toll* mutant ATAC-seq (3 h, 4 h and 5 h AEL), H3K27ac and CBP ChIP-seq (2-4 h AEL) and PRO-seq (3 h) alongside Dl ChIP-nexus (2-4 AEL wild-type embryos) ^41^ and zen (2-4 h AEL *gd^7^* embryos) and twi ChIP-seq ^13,14^ at *dpp, ind* and *twi*. The genomic positions of DV enhancers and promoters are denoted. **g)** Genome browser closeups of *Toll* mutant H3K27ac and CBP signal at the *twi* promoter and enhancer. **h)** Boxplots showing the fold change (log_2_) in enhancer RNA (eRNA) activity measured from *Toll* mutant PRO-seq at DV and non-DV enhancers. **i)** Metagene profiles of *Toll* mutant Pol II (Rpb3) CUT&Tag (2-4 h AEL) and PRO-seq (3 h) signal (RPKM) at DV enhancers (± 5 kb of CBP) and promoters (± 5 kb of TSS). Comparisons of the enrichment at DV enhancer and promoter gene classes between *Toll* mutants are from the Wilcoxon signed-rank test.

We assigned genomic regions with differential occupancy and accessibility to target genes within the same topologically associated domain (TAD), and identified 176 putative DV enhancers linked to 107 promoters (Fig. S2a,b and Table S4). Most genes were associated with one or two DV enhancers, suggesting that our approach discerns critical regulatory sequences, but a few genes had multiple enhancers (Fig. S2c). Examining the distribution of enhancer-TSS genomic distances revealed a class of promoter-proximal enhancers, but the majority of enhancers (85%) were distal (> 700 bp) from their targets (Fig. S2d). DV enhancers showed a characteristic pattern of H3K27ac flanking the central maxima of CBP enrichment and region of accessible chromatin (Fig. S2e), likely reflects CBP recruitment by DNA-binding TFs (Fig. S2e,f). We validated our enhancer identification strategy by examining overlapping genomic regions tested in a high-throughput transgenic reporter-gene assay ^32^, for which we observed enrichment of annotation terms associated with dorsal ectoderm expression for *gd^7^* enhancers, ventral ectoderm for *Toll^rm9/rm10^* enhancers and mesoderm for *Toll^10B^* enhancers (Fig. S2g). Examples of regions overlapping DV enhancers tested in reporter assays that recapitulate the expected spatial expression patterns are shown in Fig. S2h ^32^. We conclude that chromatin state data is highly efficient in identifying tissue-specific enhancers.

We categorized enhancers based on the tissue of expression of their target genes and observed high tissue specificity of elevated chromatin accessibility, CBP occupancy and H3K27ac (Fig. 2b and S2i). Strikingly, a chromatin state enhancer score based on the combined tissue-specific signal for CBP, H3K27ac and ATAC-seq could accurately predict the level of expression as determined by PRO-seq (Fig. 2c, *R^2^* values 0.78, 0.62, 0.69 for dorsal ectoderm, neuroectoderm and mesoderm enhancers, respectively), and had higher predictive value when combined than individually (Fig. S2j). The chromatin state of DV promoters varied less between tissues and predicted the expression of target genes with less accuracy (Fig. 2d,e, Fig. S2j,k). In summary, the data suggests that whereas enhancer chromatin state reflects tissue-specific expression, the chromatin state at promoters is more tissue-invariant and may allow recruitment and establishment of paused Pol II to prime DV promoters for transcription in all three germ layers.

### CBP is catalytically active at enhancers but enzymatically inactive at promoters

The observation that occupancy of the CBP coactivator is far less tissue-specific at DV promoters than at enhancers indicates that it may have distinct functional roles at enhancers and promoters. The presence of CBP at promoters does not lead to H3K27ac in tissues where the gene is silent (Fig. 2f,g), indicating that the catalytic activity of CBP is modulated. Since CBP occupancy at DV enhancers correlates with transcription and H3K27ac, the data suggest that enhancer-bound CBP is catalytically active and mediates tissue-specific H3K27ac (e.g. close-up of the *twi* locus in Fig. 2g). Whereas CBP occupancy occurred at focused enhancer and promoter regions, tissue-specific H3K27ac spreads over larger distances covering regulatory and genic regions of DV genes (Fig. 2f and S2f), indicating that transient associations of CBP with larger genomic regions may explain the dispersed H3K27ac pattern. Our data is consistent with a model where promoter-bound CBP supports Pol II recruitment and pausing in an enzymatically-independent manner ^33^, whereas catalytic CBP activity at enhancers is critical for tissue-specific histone acetylation and release from pausing.

In mammals, non-coding transcription is a predictive marker of active enhancers and enhancer RNAs (eRNAs) can allosterically activate the HAT activity of p300/CBP ^34–36^ and are implicated in supporting the transition of paused Pol II into elongation ^37^. *Drosophila* eRNA transcription also correlates with enhancer activity ^38^, but direct comparisons of eRNA levels between the same enhancer in active and inactive cellular contexts are lacking. From the PRO-seq signal at intergenic enhancers and the non-coding strand of genic enhancers, we detected eRNAs that were more abundant in the tissue where the target gene was expressed (Fig. 2h). For example, at the intronic *dpp* E1 enhancer we detected an eRNA with strong antisense transcription specific to *gd^7^* embryos (Fig. S2l). It is possible that these eRNAs contribute to activating CBP catalytic activity at enhancers ^35^.

*Drosophila* eRNAs are not as abundant as in mammals, but interestingly, Pol II CUT&Tag enrichment at DV enhancers was strong, whereas the PRO-seq signal was low compared to promoters (Fig. 2i). It therefore appears that Pol II is efficiently recruited to both promoters and enhancers, but that Pol II engages in transcription to a lesser extent at enhancers. This suggests that features specific to enhancers and promoters are involved in establishing transcriptionally engaged Pol II at a post-recruitment step.

Overall, we show that by integrating genome-wide data providing multiple indicators of chromatin state with transcription, key tissue-specific enhancers can be accurately identified. The data indicate that an active chromatin state is established by tissue-specific recruitment of CBP to enhancers, which leads to histone acetylation across DV gene loci. The finding that the chromatin state of promoter regions is less tissue-specific may reflect the uniform recruitment of paused Pol II across tissues and indicates that tissue-specific signals from the enhancer chromatin state trigger the release of paused Pol II into elongation.

### Tissue-specific transcription factors are enriched at DV enhancers

To identify transcription factors (TFs) involved in recruiting CBP and H3K27ac to DV genes, we performed a *de novo* motif analysis of DV enhancers using the MEME suite ^39^. A motif for Mad, the Smad protein that transduces Dpp signalling, was enriched in dorsal ectoderm enhancers, whereas Dl motifs were enriched in neuroectoderm and mesoderm enhancers (Table S5). We then plotted the enrichment of known motifs from the JASPAR database ^40^ at DV enhancers, which revealed that Zelda (Zld) and Brinker (Brk) motifs were the most strongly enriched in dorsal ectoderm enhancers, Snail (Sna) and Dl in neuroectoderm, and Su(H) (Suppressor of Hairless) and Twist (Twi) motifs in mesoderm enhancers (Fig. S3a-b and Table S6). We also examined published ChIP-nexus data for Dl ^41^ and ChIP-seq data for several tissue-specific TFs ^13,14^, which showed that Mad and Zerknüllt (Zen) occupancy is common in dorsal ectoderm enhancers, Dl in neuroectoderm enhancers, whereas Twi binding is enriched in mesoderm enhancers (Fig. S3c). Locus-specific occupancy is also evident in the browser snapshots of *dpp, ind* and *twi* (Fig. 2f). Together, these analyses show that tissue-specific transcriptional activators (e.g. Mad, Zen, Twi) and repressors (Brk and Sna) have motifs in and bind to the identified DV enhancers. They may contribute to differential CBP and H3K27ac recruitment, but must act after or in parallel to Dl, since the genes encoding these TFs are themselves targets of the Dl gradient.

### DV transcription occurs within the context of a tissue-invariant chromatin conformation

Early *Drosophila* embryogenesis involves the rapid formation of an elaborate 3D chromatin organization characterized by the establishment of TADs and the formation of enhancer-promoter loops ^42,43^. Although TAD formation coincides with ZGA, it occurs independently of transcription, is tissue invariant and gene expression is largely unaltered by major disruptions of chromosome topology ^7,42,44^. Enhancer-promoter loops are also maintained across tissues in the early embryo ^7,8,42,43,45^, so although these loops are important for positioning enhancers and promoters in proximity to each other, additional regulatory components are required to drive tissue-specific expression. Consistently, despite the major differences in chromatin state and transcription, the genome organization of the DV-regulated genes *dpp*, *ind* and *twi* appear largely tissue invariant between *Toll* mutants (Fig. S3d) ^7^.

### Tissue-specific P-TEFb recruitment releases Pol II into productive elongation at DV genes

Since both chromatin conformation as well as the chromatin state at promoters is largely tissue invariant and may reflect the uniform recruitment of paused Pol II across tissues, signals from the enhancer chromatin state may trigger the release of paused Pol II into elongation. A critical step in the release of paused Pol II is the phosphorylation of negative elongation factors and the Pol II C-terminal domain (CTD) by the P-TEFb kinase, consisting of CDK9 and Cyclin T (CycT) (Fig. 3a) ^46,47^. To investigate if tissue-specific activity of P-TEFb at DV genes is regulated by differential recruitment or enzymatic activation, we performed CycT and CDK9 CUT&Tag in 2-4 h *Toll* mutant embryos. This revealed that P-TEFb occupancy is more strongly associated with dorsal ectoderm promoters in *gd^7^* embryos, neuroectoderm promoters in *Toll^rm9/rm10^* embryos, and mesoderm promoters in *Toll^10B^* embryos (Fig. 3b and S4a). We validated this result by ChIP-qPCR, showing tissue-specific CycT enrichment at DV promoters (Fig. S4b). Interestingly, we observed comparable levels of enrichment, and even higher tissue-specificity at DV enhancers (Fig. 3b,c and S4c). This suggests that enhancer-binding factors load the P-TEFb complex and direct it to the corresponding promoter. To test this, we investigated P-TEFb occupancy at the *Dorsocross* (*Doc*) locus that consists of three genes (*Doc1, Doc2 and Doc3*) and five enhancers (Fig. 3d). CycT was highly enriched at both enhancers and promoters in the dorsal ectoderm (*gd^7^* embryos) compared to the other tissue-types. We then examined CycT occupancy in embryos homozygous for a deletion of the *Doc* E1 enhancer ^8^. Removal of this single enhancer marginally reduced expression of the *Doc* genes and had minimal effects on the chromatin state of the locus (Fig. S4d,e), reflecting functional redundancy with the intact enhancers that maintain promoter contacts ^8^. Nevertheless, by ChIP-qPCR we could detect a reduction in the occupancy of CycT at the *Doc* promoters in embryos lacking the E1 enhancer (Fig. 3e), indicating that enhancers modulate loading of P-TEFb to promoters.

**Figure 3.**
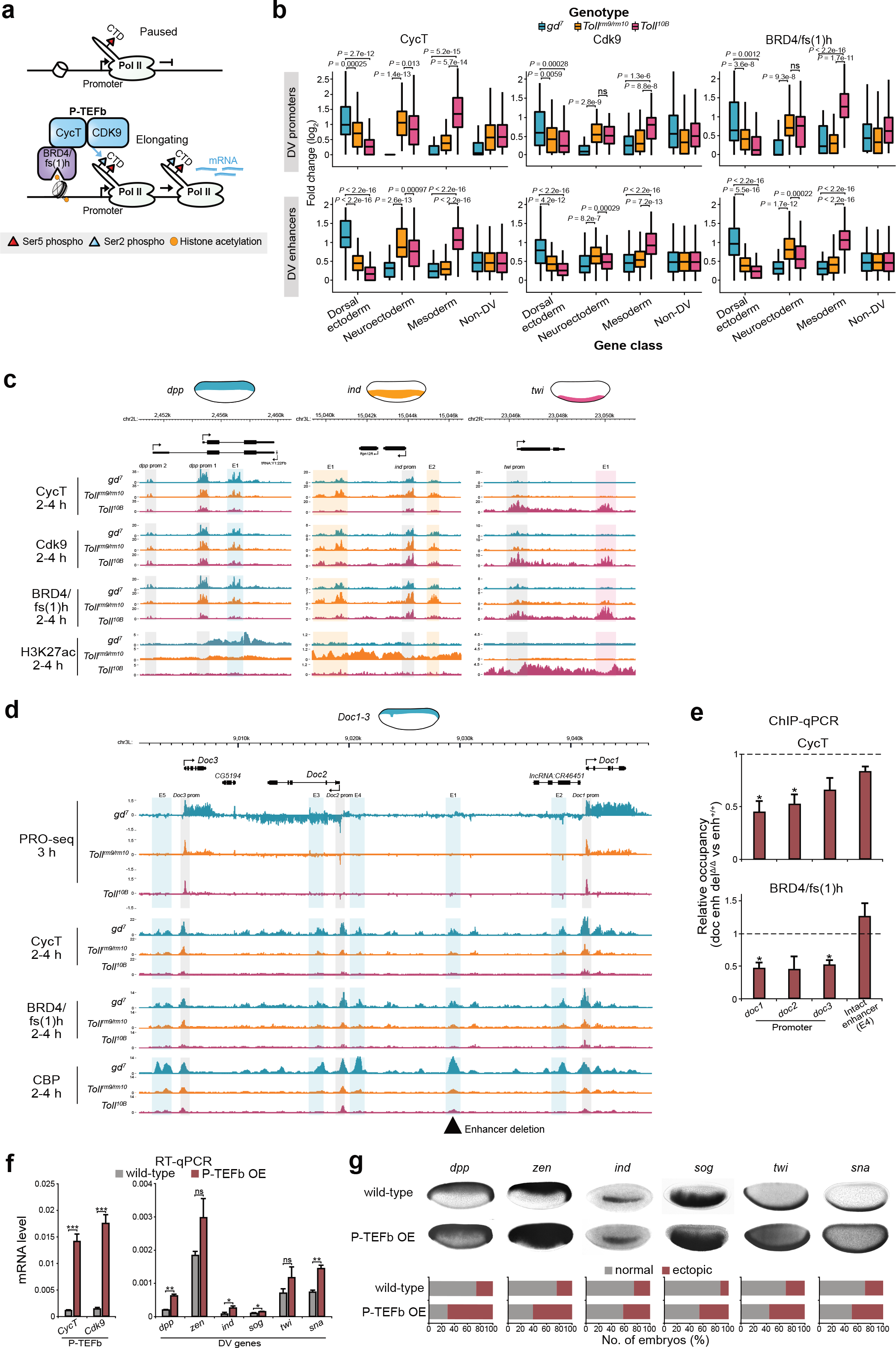
Tissue-specific P-TEFb recruitment is associated with the release of paused Pol II into productive elongation at DV promoters. **a)** Schematic of P-TEFb (composed of CycT and CDK9 subunits) mediated release of promoter-proximal paused Pol II into productive elongation. P-TEFb phosphorylates serine 2 of the Pol II carboxyl-terminal domain (CTD) to stimulate elongation. BRD4/fs(1)h binds to acetylated histones and helps recruit P-TEFb to promoters. **b)** Boxplots showing the fold change (log_2_) in enrichment of CycT, Cdk9 and BRD4/fs(1)h from CUT&Tag in *Toll* mutant embryos at DV promoters and enhancers. Comparisons of the enrichment at DV enhancer and promoter gene classes between *Toll* mutants are from the Wilcoxon signed-rank test. **c)** Genome browser shots of *Toll* mutant CycT, Cdk9 and BRD4/fs(1)h CUT&Tag and H3K27ac ChIP-seq at *dpp*, *ind* and *twi*. **d)** Genome browser shot of *Toll* mutant PRO-seq, CBP ChIP-seq and CycT and BRD4/fs(1)h CUT&Tag at the *doc* locus. The position of the *doc* E1 enhancer deletion ^8^ is denoted. **e)** ChIP-qPCR showing the enrichment of CycT and BRD4/fs(1)h at the promoters of *doc1*, *doc2* and *doc3* in doc enh del^Δ/Δ^ embryos (2-4 h AEL) relative to enh^+/+^ embryos (*n* = 3-4). Relative occupancy is also shown at the intact *doc* enhancer (E4). Error bars show SEM. Significant differences in occupancy (two tailed, unpaired t-test) are indicated by asterisks (* = *P* < 0.05). **f)** RT-qPCR quantification of *CycT*, *Cdk9* and DV regulated genes (*dpp*, *zen*, *ind*, *sog*, *twi* and *sna*) mRNA levels (relative to 28S rRNA) in wild-type embryos and P-TEFb maternally overexpressed (OE) embryos. Error bars show SEM. Significant differences in mRNA (two tailed, unpaired t-test) are indicated by asterisks (* = *P* < 0.05, ** = *P* < 0.01, *** = *P* < 0.001). **g)** (top) Images of whole mount *in situ* hybridization in wild-type and P-TEFb OE mutant embryos (2-4 h AEL) with probes hybridizing to mRNAs of the DV regulated genes in **f** and (bottom) quantification of the proportion of embryos with normal or ectopic staining for each probe. The number of embryos sampled are detailed in the methods.

One factor that has been implicated in P-TEFb recruitment is the tandem bromo- and extra-terminal domain (BET) protein BRD4, known as female sterile (1) homeotic (fs(1)h) in *Drosophila* ^48,49^. We performed BRD4/fs(1)h CUT&Tag and found that it is also more strongly associated with DV promoters and enhancers in the tissue of target gene expression (Fig. 3b-d, S4a and c), and that occupancy at the *Doc* promoters was reduced in the absence of the E1 enhancer (Fig. 3e). Although BRD4/fs(1)h can recognize acetylated histones through its bromodomains, occupancy was restricted to enhancers and promoters and did not overlap the more distributed H3K27ac pattern (Fig. 3c and S4a). This indicates that other histone modifications or factors binding accessible chromatin at enhancers may be more important for BRD4/fs(1)h recruitment than H3K27ac.

Tissue-specific enrichment of P-TEFb suggests that this kinase could be limiting for transcription in non-expressing tissues. We therefore over-expressed Cdk9 and CycT in early embryos with the maternal tub-Gal4 driver, leading to more than 10-fold increased expression in embryos (Fig. 3f). Although occupancy of P-TEFb did not increase at tested promoters according to CycT ChIP-qPCR (Fig. S4f), expression of DV genes was elevated (Fig. 3f). Interestingly, the number of embryos with DV expression detected outside the normal expression domain was significantly increased by P-TEFb over-expression for all DV genes examined by whole-mount *in situ* hybridization (Fig. 3g and S4g). Ectopic expression may result from titration of negative regulators of P-TEFb, such as the 7SK snRNP that sequesters and inactivates the kinase ^50^, since promoter occupancy did not change upon P-TEFb overexpression. Consistent with this notion, the frequency of ectopic expression correlated with the level of CycT at gene promoters in non-expressing tissues (*R* = 0.72, Fig. S4h).

Together, the results suggest that both regulated recruitment of P-TEFb as well as relief from inhibition may be important for tissue-specific release of Pol II from promoter-proximal pausing. Our data show that BRD4/fs(1)h and P-TEFb are enriched in the tissue where the genes are expressed and suggest that differential recruitment of these factors leads to pause release, but indicate that activation of P-TEFb kinase activity is also necessary.

### Repressors block the release of paused Poll II into elongation and exclude H3K27ac or induce Polycomb-mediated H3K27me3

To decipher if tissue-specific control of Pol II pausing requires active repression in non-expressing cells, we compared the chromatin state at genes regulated by the Dl and Snail repressors (Fig. 4a). Dl is converted to a repressor when its binding sites are flanked by AT-rich elements that recruit Capicua (Cic) and the co-repressor Groucho, resulting in long-range repression to delimit the ventral boundary of dorsal ectoderm specific genes ^51,52^. In the mesoderm, Snail (Sna) works as a short-range repressor by recruiting the CtBP and Ebi co-repressors to shut down neuroectoderm-specific enhancers ^15,53,54^. Publicly available Dl ChIP-nexus ^41^ and Sna ChIP-seq ^13,14^ data identified peaks in 5-33% of DV enhancers (Fig. S5a), allowing us to identify 13 putative Sna-target genes and 6 Dl-target genes (Fig. 4).

**Figure 4.**
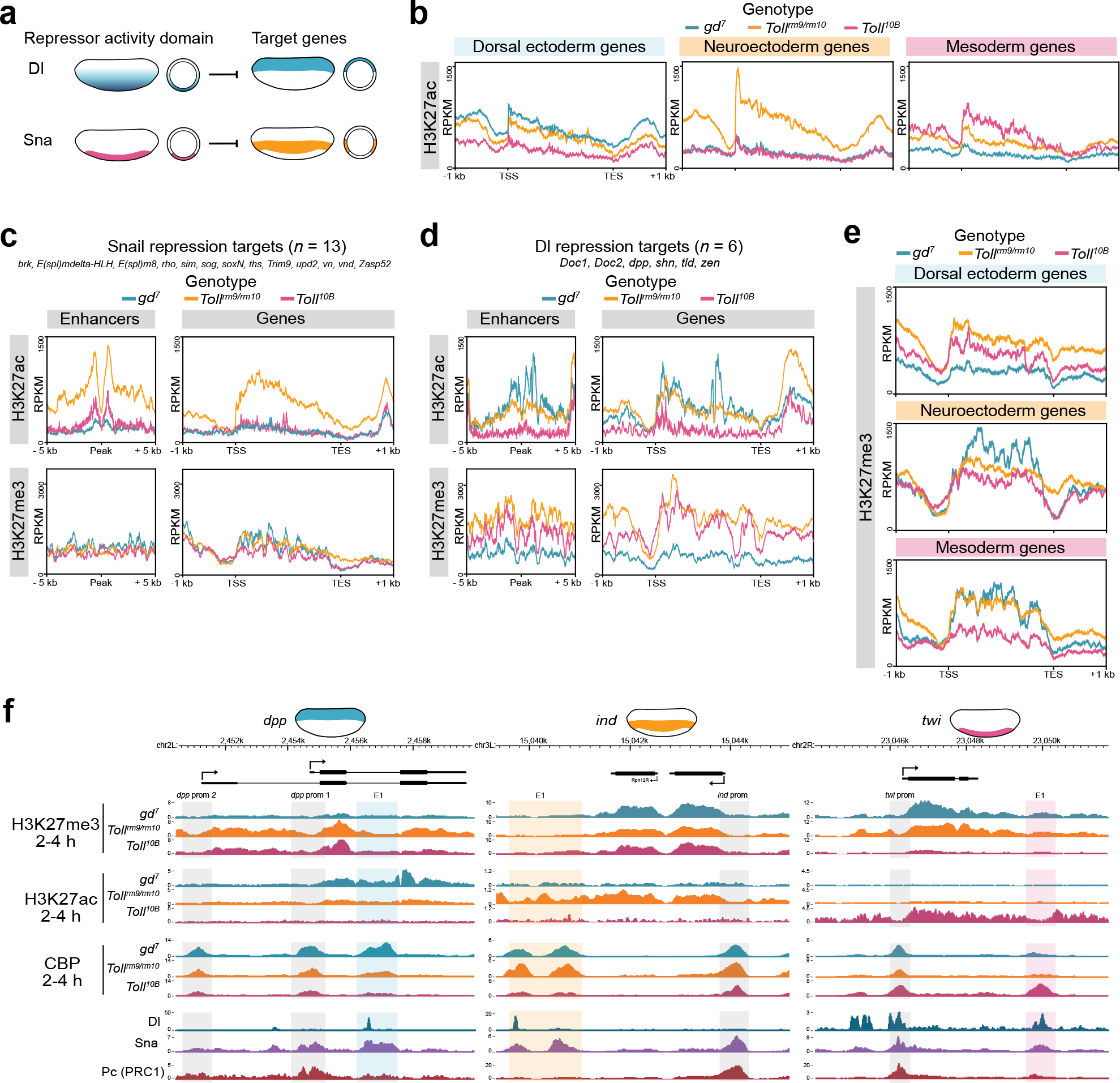
Repressors prevent the release of paused Pol II at DV promoters by excluding H3K27ac and recruiting Polycomb-mediated H3K27me3. **a)** Schematic of the repressor activity domains and target genes of Dl and Sna-mediated repression. **b)** Metagene plots of *Toll* mutant H3K27ac ChIP-seq (2-4 h AEL) signal at DV regulated genes. **c-d)** Metagene plots of *Toll* mutant H3K27ac and H3K27me3 ChIP-seq at enhancers and gene bodies of Sna **(c)** and Dl **(d)** repressor targets. **e)** Metagene plots of *Toll* mutant H3K27me3 ChIP-seq signal at DV regulated genes. **f)** Genome browser shots of *Toll* mutant H3K27me3, H3K27ac and CBP ChIP-seq signal alongside Dl ChIP-nexus and Sna and Pc ChIP-seq from wild-type (2-4 h AEL) embryos at *dpp*, *ind* and *twi*.

We found that the Sna repressor did not prevent occupancy of the Dl activator or induce chromatin compaction in the mesoderm (Fig. S5b,c). Instead, prevention of H3K27ac at DV loci appears to be a major target of Sna-mediated repression (Figs. 2b, 4b,c). This suggests that Sna quenches the Dl activator in the mesoderm by preventing CBP-mediated H3K27ac. By contrast, when Dl acts as a repressor, it does not block H3K27ac at its targets in the neuroectoderm, although these genes are hypoacetylated in the mesoderm (Fig. 4d). Instead, the Polycomb-catalyzed mark H3K27me3 accumulates at Dl-repressed targets in both the neuroectoderm and mesoderm (Fig. 4d), indicating that Polycomb silencing is an important enforcer of Dl-mediated repression. However, Sna-targets did not accumulate H3K27me3 in the mesoderm (Fig. 4c), consistent with the notion that Sna represses transcription by a different mechanism.

H3K27me3, which anti-correlates with DV gene activation ^13^, accumulates across genomic regions that encompass the gene bodies of DV genes in a tissue-specific manner (Fig. 4e, f). However, Polycomb group proteins (PcGs) have been shown to interfere with Pol II initiation and elongation checkpoints ^55–57^. ChIP-seq data for the Polycomb Repressive Complex 1 (PRC1) component Polycomb (Pc) in 2-4 h AEL wildtype embryos ^13,14^ detected Pc enrichment specifically at DV promoters and not at enhancers (Fig. S5d,e). We found that 39% of the DV promoters, but only 4% of enhancers, overlapped known *Drosophila* Polycomb Response Elements (PREs) (Fig. S5f) ^58^. The strong promoter bias of Pc occupancy suggests that silencing may also involve impeding Pol II activity post-recruitment at promoters. Consistent with this is the retention of similar levels of paused Pol II at DV promoters across tissues, suggesting that DV repressors inhibit the release of paused Pol II ^59^. Together, our data show that whereas Dl-mediated repression is accompanied by PcG silencing and H3K27me3, repression by Sna involves prevention of H3K27ac without induction of H3K27me3, but both mechanisms impair the release of paused Pol II into elongation.

### DV enhancers and promoters are temporally primed by pioneer factors for increased accessibility prior to induction of DV transcription

We next aimed to complement our tissue-resolved map of the activity of DV enhancers and promoters by temporally resolving chromatin and transcriptional state dynamics during DV patterning (Fig. 5a). We plotted the temporal dynamics of chromatin accessibility at DV enhancers and promoters using ATAC-seq data from wild-type embryos through nuclear cycles 11-13, immediately preceding ZGA ^60^. Since the Dl gradient response gradually appears between nuclear cycles (nc) 12-14, we expect chromatin accessibility to be largely uniform across cells in wild-type embryos during nc 11-13. We found that both DV enhancers and promoters are more accessible than shuffled sites representative of the genomic background prior to the initiation of DV gene transcription (Fig. 5b).

**Figure 5.**
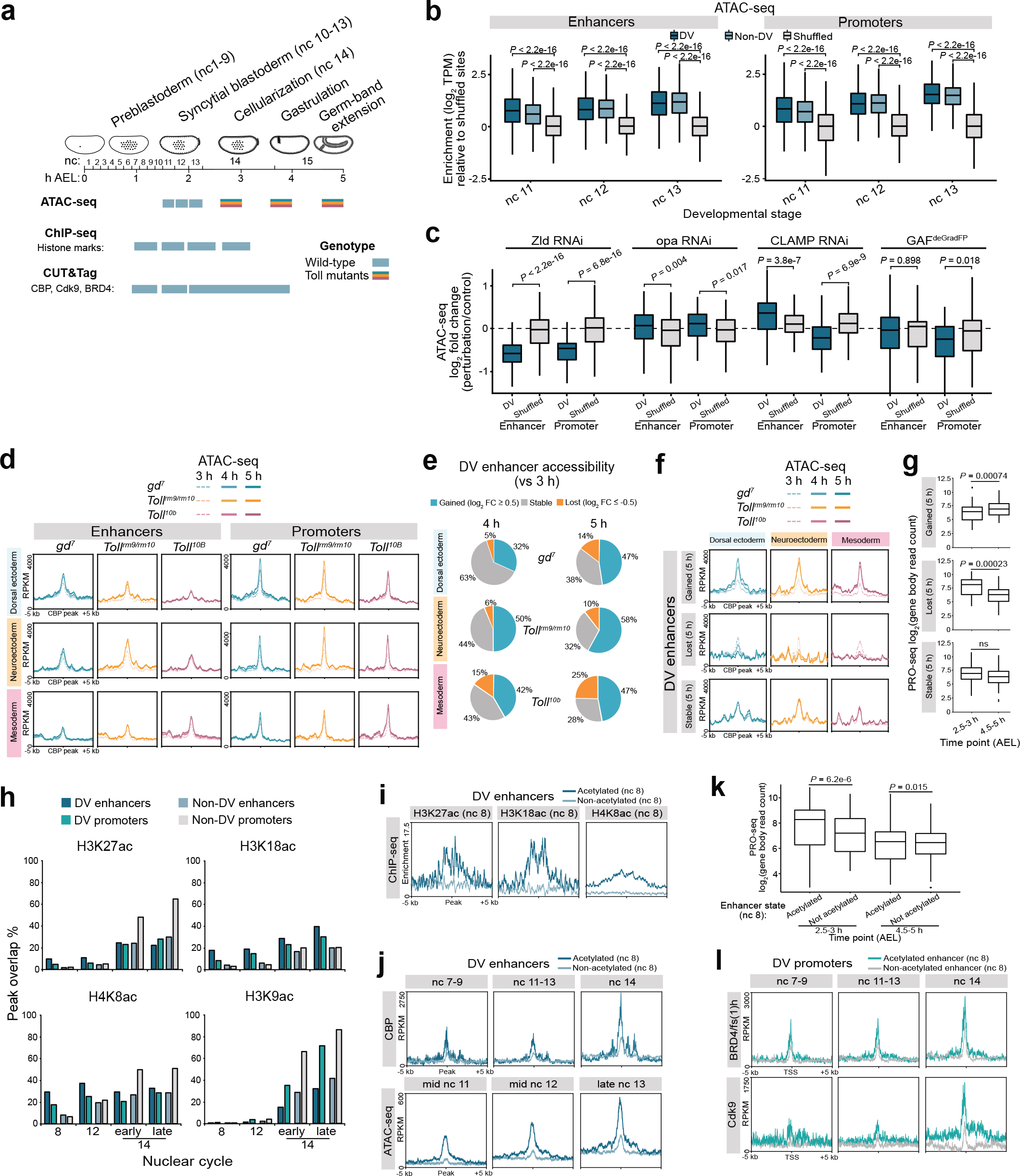
DV regulated enhancers are temporally primed by increased chromatin accessibility and CBP-mediated histone acetylation prior to the commencement of DV transcription. **a)** Schematic of the developmental stages profiled by ATAC-seq, ChIP-seq and CUT&Tag. The nuclear cycles (nc) and hours after egg laying (hAEL) are indicated. **b)** Boxplots of ATAC-seq enrichment (log_2_ TPM) at DV and non-DV enhancers and promoters, relative to shuffled genomic regions from wild-type embryos at nc 11, 12 and 13 ^60^. **c)** Boxplots showing the log_2_ fold change (perturbation/control) in ATAC-seq signal at DV and shuffled enhancers and promoters after maternal RNAi depletion of Zld and opa ^63^, CLAMP ^64^ and zygotic GAF^deGradFP 65^. *P*-values (Wilcoxon rank-sum test) show significant differences in accessibility compared to shuffled sites. **d)** Metagene plots of *Toll* mutant embryo (3 h, 4 h and 5 h AEL) ATAC-seq signal (RPKM) at DV enhancers and promoters partitioned by the tissue of target gene activity. **e)** Proportion (%) of dorsal ectoderm, neuroectoderm and mesoderm enhancers that gained (log_2_ fold change (FC) ≥ 0.5), lost (log_2_ FC ≤ −0.5) or maintained stable chromatin accessibility (ATAC-seq) in 4 h and 5 h AEL embryos relative to 3 h. The accessibility was measured from *gd^7^* at dorsal ectoderm enhancers, *Toll^rm9/rm10^* at neuroectoderm enhancers and *Toll^10B^* at mesoderm enhancers. **f)** Metagene plots of *Toll* mutant ATAC-seq signal (3 h, 4 h and 5 h AEL) at DV enhancers partitioned by the tissue of target gene activity and the change of accessibility (5 h relative to 3 h AEL). **g)** Boxplots of early (2.5-3 h) and late (4.5-5 h) *Toll* mutant PRO-seq gene body expression (log_2_ read count) of DV genes associated to enhancers with gained, lost or stable accessibility (5 h vs 3 h). Expression was measured for genes in the tissue mutant of activity. *P*-values (Wilcoxon rank-sum test) show significant differences in expression (2.5-3 h vs 4.5-5 h). **h)** Overlap (%) of DV and non-DV enhancers and promoters with ChIP-seq peaks (nc 8, 12, 14 (early and late)) called for the p300/CBP-mediated histone acetylation marks (H3K27ac, H3K18ac and H4K8ac) and the non-p300/CBP mark H3K9ac ^66^. **i-j)** Metagene plots of **(i)** CBP-catalyzed histone marks from nc 8 ChIP-seq and **(j)** CBP CUT&Tag and ATAC-seq enrichment at DV enhancers acetylated or non-acetylated at nc 8. **k)** Boxplots of 2.5-3 h and 4.5-5 h (AEL) PRO-seq gene body read counts (log_2_) for DV genes linked to enhancers acetylated or not acetylated at nc 8. For each gene, the PRO-seq signal was taken from the *Toll* mutant of expression. *P*-values are from the Wilcoxon rank-sum test. **l)** Metagene plots of BRD4/fs(1)h and Cdk9 CUT&Tag signal from nc 7-9, 11-13 and 14 wild-type embryos at the promoters of DV genes linked to enhancers acetylated or non-acetylated at nc 8.

Consistent with the early priming of DV regulatory elements, the pioneer factor Zld, which has been shown to potentiate Dl activity at DV enhancers ^61^, is highly enriched at DV enhancers and promoters already in nc 8 embryos (Fig. S6a,b) ^62^. Alongside Zld, three factors with pioneer-like activities in the early embryo have been identified, Odd-paired (Opa) ^63^, CLAMP ^64^ and GAGA-factor (GAF, also known as Trithorax-like, Trl) ^65^. We found that Opa and CLAMP occupy both DV enhancers and promoters, whereas GAF favours DV promoters (Fig. S6c). We analyzed published ATAC-seq data from embryos where each pioneer factor had been perturbed individually (Fig. 5c). We observed a small loss of accessibility at DV promoters upon *GAF* inactivation, an unexpected slight increase in enhancer accessibility in *CLAMP* RNAi embryos, virtually no effect of *opa* RNAi, and a more pronounced loss at both DV enhancers and promoters in *zld* RNAi embryos ^63–65^ (Fig. 5c). This is consistent with earlier work demonstrating a function for Zld in expression and accessibility of DV genes ^61,62^.

### Temporal changes to enhancer accessibility correlate with variations in DV expression

To explore spatio-temporal accessibility dynamics during the induction of DV responsive transcription, we analyzed our *Toll* mutant ATAC-seq data from three time points covering early, intermediate and late phases of ZGA (3 h, 4 h and 5 h AEL, respectively) (Fig. 5a). ATAC-seq revealed that tissue-specific differences in the accessibility of DV enhancers were augmented from 3 h to 5 h (Fig. 5d). DV promoters, although less tissue-specific in accessibility, also gained accessibility over time across tissues (Fig. 5d). We quantified changes in accessibility across the time course for DV enhancers specifically in the tissue of target gene activity, to identify enhancers that gained (log_2_ fold change ≥ 0.5), lost (log_2_ fold change ≤ – 0.5) or retained stable accessibility (Fig. 5e). While the majority of enhancers gained accessibility or remained stably open over time in the tissue of expression, some lost accessibility (Fig. 5e,f). Measuring the PRO-seq gene body expression at early (2.5-3 h) and late (4.5-5 h) phases of DV-responsive transcription revealed corresponding effects on transcription (Fig. 5g).

The closing down of specific enhancers may indicate transfers of regulatory control between enhancers that drive different spatio-temporal expression patterns of the same gene. The locus encoding the dorsal ectoderm-expressed gene *schnurri* (*shn*) exemplifies how chromatin accessibility changes at enhancers can correspond to their spatio-temporal activities, while promoter accessibility can be maintained or gained across tissues (Fig. S6d-f). The *shn* E1 enhancer is primed by chromatin accessibility through nc 11-13, and as a result has accessibility in all the tissue mutants at the start of nc 14 (3 h) (Fig. S6e,f). The E1 enhancer closes down in *Toll^rm9/rm10^* and *Toll^10B^* embryos at 4 h, and in *gd*^7^ embryos at 5 h. Decommissioning of E1 occurs concomitantly with a gain in accessibility for the upstream E2 enhancer specifically in *gd*^7^ embryos from 4 h onwards, suggesting regulatory control of *shn* is transferred from E1 to E2 as development proceeds (Fig. S6e,f). In contrast, accessibility at the *shn* promoter increases from 3 h to 5 h in a largely tissue invariant manner. In support of E1 and E2 driving early and late *shn* expression, reporter gene activities driven by fragments overlapping E1 and E2 have distinct spatial and temporal patterns that recapitulate the early and later embryonic expression patterns of *shn*, respectively (Fig. S6f) ^32^.

The data suggest that DV enhancers become accessible prior to DV gene transcription with the help of Zelda, and that dynamic alterations in accessibility correlate with spatial and temporal changes in transcription. By contrast, DV promoters maintain more stable accessibility across tissues consistent with the retention of paused Pol II.

### CBP-mediated acetylation primes a subset of DV enhancers for rapid induction of tissue-specific transcription

To investigate if the temporal priming of chromatin accessibility at DV enhancers and promoters is accompanied by changes in histone modifications, we examined spike-in normalized ChIP-seq data for a wide range of histone marks from nc 8, 12, 14a (early) and 14c (late) wild-type embryos (Fig. 5h and Fig. S6g) ^66^. This showed that the CBP-catalyzed marks H3K27ac, H3K18ac and H4K8ac gradually accumulated at enhancers and promoters, with enrichment elevated relative to shuffled regions already by nc 8 (Fig. S6g). By contrast, deposition of non-CBP catalyzed H3K9ac, and methylation of H3K4 (H3K4me1/me3) occurred co-transcriptionally at nc 14 (Fig. 5h and Fig. S6g). Interestingly, a greater proportion of DV enhancers than non-DV enhancers were marked with H3K27ac, H3K18ac and H4K8ac prior to ZGA, but by nc 14 the overlap was similar between DV and non-DV enhancers (Fig. 5h). We determined the overlap of the enhancers with acetylation over time (Fig. S6h), and identified 48 DV enhancers with any CBP catalyzed acetylation already present at nc 8 (Fig. 5i). Of these, 96% overlap Zld ChIP-seq peaks from the same stage, compared to 46% of the non-acetylated DV enhancers (Fig. S6i) ^66^.

The deposition of histone acetylation at a subset of DV enhancers prior to ZGA suggests that CBP is recruited to chromatin before DV transcription commences. To test this, we performed CUT&Tag on hand-sorted nc 7-9, 11-13, and 14 embryos, which demonstrated that CBP was enriched at DV enhancers and promoters relative to shuffled genomic regions already at nc 7-9 (Fig. S6j). The Zld-bound early acetylated DV enhancers were more enriched for CBP and had markedly higher accessibility than non-acetylated enhancers across the pre-ZGA nuclear cycles (Fig. 5j).

To assess whether the early establishment of an active chromatin state at a subset of DV enhancers influenced target genes, we examined the chromatin state at DV promoters. Promoters linked to the early acetylated enhancers were also more enriched for histone acetylation than promoters linked to non-acetylated enhancers, were more accessible, and had stronger CBP enrichment (Fig. S6k,l). To see whether the early established active chromatin states influenced DV transcription, we plotted the PRO-seq gene body expression for target genes from the tissue mutant of expression at early (2.5-3 h) and late (4.5-5 h) stages (Fig. 5k). PRO-seq revealed that DV genes with early established active enhancer and promoter chromatin states established stronger tissue-specific transcription at the beginning of nc 14 (2.5-3 h AEL). Thus, our data suggest that a subset of DV enhancers are primed by Zld for rapid establishment of an active chromatin state defined by elevated chromatin accessibility, recruitment of CBP and enrichment of CBP-catalyzed histone acetylations, and that this results in rapid induction of tissue-specific transcription.

### Strong P-TEFb enrichment at DV promoters is not observed until gene expression is initiated

Since DV genes are paused but not expressed in naïve embryos, we examined when P-TEFb and BRD4/fs(1)h became associated with these genes. We performed CUT&Tag for CDK9 and BRD4/fs(1)h on nc 7-9, 11-13 and 14 embryos. We detected significant enrichment of BRD4/fs(1)h at DV enhancers and promoters, relative to shuffled genomic regions, already at nc 7-9 (Fig. S6j). The promoters of DV genes with early established enhancer and promoter chromatin states and stronger initiation of tissue-specific transcription at nc 14 also had stronger enrichment of BRD4/fs(1)h than other DV promoters across the time course (Fig. 5l). Interestingly, although weak enrichment of CDK9 was observed at nc 7-9 and 11-13 at DV promoters linked to both early acetylated and not-acetylated enhancers, strong CDK9 recruitment occurred concomitantly with the induction of expression at nc 14, with promoters linked to early acetylated enhancers having the strongest occupancy (Fig. 5l).

Taken together, the data suggest that DV enhancers are temporally primed by the pioneer factor Zld leading to an active chromatin state and BRD4/fs(1)h recruitment prior to the induction of DV-responsive transcription. However, strong loading of P-TEFb to the promoter does not occur until nc 14, which may be the trigger for the rapid release of paused Pol II and induction of tissue-specific gene expression.

### Identification of DV cell clusters from single-cell expression data

Quantitative studies have revealed that transcription is stochastic and occurs in bursts ^67^. Our results show that the DV genes are regulated by pause release, but mediation of the release of paused Pol II to produce bursts of transcription is poorly understood. We analyzed single-cell RNA-seq (scRNA-seq) data from wild-type and *Toll* mutant 2.5-3.5 h (AEL) embryos ^7^ to link these processes. Clustering of single-cell expression profiles previously identified 15 clusters representing different cell identities in the early embryo (Fig. S7a) ^7^. We performed principal component analysis (PCA) using the DV genes identified by PRO-seq on cells from the ectoderm, neural and mesoderm clusters, and used shared nearest neighbor (SNN) clustering on the first 10 principal components to assign 6 new clusters and visualized it with Uniform Manifold Approximation and Projection (UMAP) (Fig. 6a). Mapping the expression of dorsal ectoderm-, neuroectoderm- and mesoderm-specific genes in these cells showed that they distinguish different clusters of cells on the UMAP (Fig. S7b). The expression of marker genes was used to identify clusters as dorsal ectoderm (*dpp, Doc1, ush*), neuroectoderm (*ind, sog, brk*), older neural cells (*scrt, ase, nerfin-1*), mesoderm (*twi, sna*), older mesoderm or myoblasts (*Mef2, meso18E, sns, sing*) and a common cell cluster (Figs. 6a and S7c, Table S7). UMAPs from *gd^7^*, *Toll^rm9/rm10^* and *Toll^10B^* embryos revealed the absence of specific clusters in mutant embryos (Fig. 6a). Mesoderm cells were almost completely absent in *gd^7^* and *Toll^rm9/rm10^* embryos, and dorsal ectoderm cells were mostly depleted from *Toll^rm9/rm10^* and *Toll^10B^* embryos, whereas neuroectoderm cells were largely missing from *gd^7^* but only moderately reduced in *Toll^10B^* embryos (Fig. 6a and S7d). This shows that the mutant embryos largely reflect the three presumptive germ layers, but that *Toll^10B^* embryos consist of 49% mesoderm cells and 34% cells that resemble neuroectoderm (Fig. S7d). The scRNA-seq profiles of *dpp*, *ind* and *twi* in *Toll* mutant embryos are shown in Fig. S7e.

**Figure 6.**
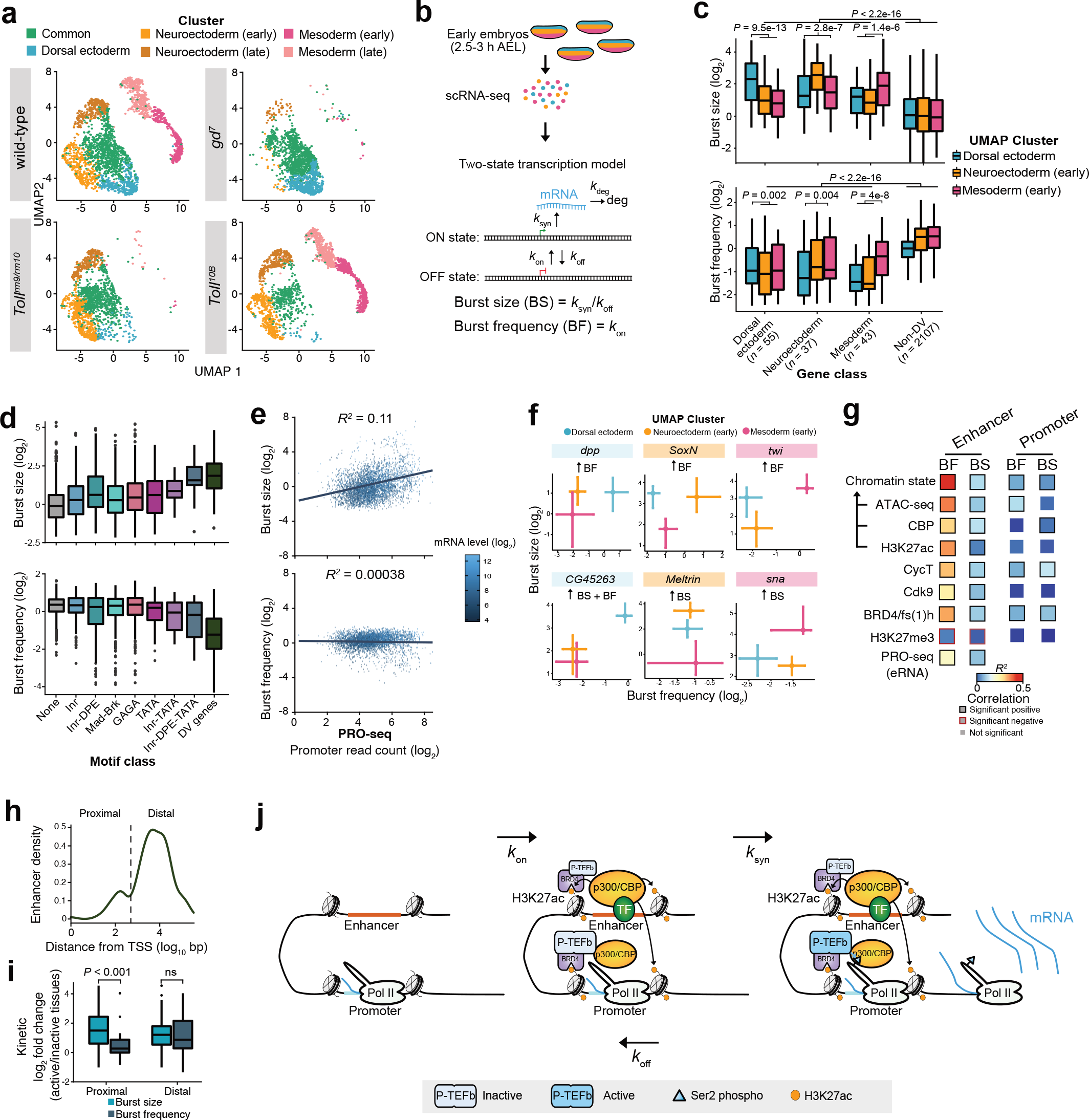
Transcriptional kinetics inferred from scRNA-seq data show that DV genes have a high burst size and are regulated in burst size or frequency. **a)** UMAP clustering of single-cell RNA-seq (scRNA-seq) from DV relevant clusters in wild-type and *Toll* mutant 2.5-3.5h embryos ^7^ based on the expression of DV genes identified by PRO-seq. **b)** Schematic of the two-state transcriptional model used for transcriptome-wide inference of burst kinetics from scRNA-seq ^68^. **c)** Boxplots showing the burst size and frequency (log_2_) of DV genes classified by the tissue of expression in DV relevant UMAP clusters from wild-type scRNA-seq. **d)** Boxplots of the burst size and frequency of genes classified by the presence of *de novo* identified promoter motifs and compared to all DV genes. **e)** Correlations between transcriptional kinetics and PRO-seq promoter read counts (log_2_). The mRNA level (log_2_ TPM) of genes is denoted. **f)** Plots showing the transcriptional kinetics of individual DV genes (*dpp*, *CG45263*, *SoxN*, *Meltrin*, *twi* and *sna*) across DV relevant UMAP clusters. Error bars show the 95% confidence intervals. Genes with statistically significant increases in bursting kinetics in the cluster of expression relative to the OFF clusters are denoted. **g)** Heatmap showing the coefficient of determination (*R^2^*) between the enrichment of various genomic datasets at DV enhancers and promoters compared to burst frequency (BF) or size (BS). Comparisons with significant positive and negative correlations are denoted by boxes. **h)** Distribution of DV enhancer density in respect of genomic distance from the TSS of target genes with inferred kinetics (log_10_ bp). Enhancers with a distance ≤ 700 bp from the TSS were classified as proximal (*n* = 22) and above this threshold defined as distal (*n* = 115). **i)** Boxplots showing the fold change (log_2_) in transcriptional kinetics between the ON tissue and mean of the OFF tissues for genes regulated by proximal and distal enhancers. **j)** Schematic model of DV gene activation.

### Transcriptome-wide inference of burst kinetics in different cell types reveals that DV genes have high burst size capacities and constrained burst frequencies

The scRNA-seq data from wild-type embryos was used to infer transcriptional burst kinetics based on a two-state model of transcription ^68^ (Fig 6b). The two-state model consists of four parameters that may accommodate different transcriptional kinetics. The rate of transition to the active state, *k*_on_; the rate of transition to the inactive state, *k*_off_; the rate of transcription in the active state, *k*_syn_; and the mRNA degradation rate, *k*_deg_. Here, we mainly characterized bursting by the burst frequency (*k*_on_; in units of mean mRNA degradation rate) and burst size (mean number of transcripts produced per active burst; *k*_syn_/*k*_off_). We modelled gene expression using bootstrapped maximum likelihood inferences to obtain estimates and confidence intervals on burst frequency and size ^68^, and removed genes with no or low burst size (non-expressed) and uncertain kinetic parameters. Burst kinetics were determined for a total of 2232 genes, including 125 DV genes, in cells of the dorsal ectoderm, neuroectoderm and mesoderm clusters (Table S8), and the kinetic values inferred were highly concordant between two different wild-type lines (*w^1118^* and *PCNA:eGFP*, Fig. S8a). The analysis revealed that DV genes have high burst sizes and low burst frequencies compared to non-DV genes, suggesting that they fire infrequently but produce many transcripts per burst (Fig 6c and S8b). Since high burst size has previously been associated with the occurrence of certain core promoter motifs _68_, we plotted the burst sizes and frequencies of genes associated with no motifs, with individual motifs or with a combination of promoter motifs (Fig. 6d). This showed that genes associated with Inr, DPE and TATA had a high burst size but low burst frequency. Since these motifs are overrepresented in the DV genes (Fig. S1j-m), it may partly explain their high burst size capacity. We also plotted the burst size and frequency relative to the level of Pol II promoter-proximal pausing genome-wide (Fig. 6e). We noted a correlation between pausing and burst size, but not burst frequency. Pausing correlated better with burst size than burst frequency also for DV genes but the correlation was weaker, likely due to the small sample size (Fig. S8c). Thus, an enrichment of core promoter motifs and high pausing may explain the high burst size of DV genes.

By comparing the burst kinetics inferred in the dorsal ectoderm, neuroectoderm (early) and mesoderm (early) cell clusters, we were able to measure changes in DV gene burst sizes and frequencies between cells where these genes are active or inactive (Fig. 6c). Comparison between the clusters showed that both burst size and burst frequency were significantly higher for DV genes in the cluster of expression (Fig. 6c). To explore whether the relative contributions of burst size and frequency parameters vary between genes, we plotted the burst size and frequency with confidence intervals for individual DV genes in the three clusters (Fig. 6f, Table S8). This revealed that DV genes have different dependencies on burst size and frequency changes during bursts. We found that of the 47 PRO-seq identified DV genes with a significant change in one or both kinetic values, 16 significantly changed in burst frequency (e.g. *dpp, SoxN and twi*), 25 increased in burst size (e.g. *Meltrin* and *sna*) and 6 genes changed in both burst size and frequency (e.g. *CG45263*) (Fig. 6f, Fig. S8d-f). There were more dorsal ectoderm and neuroectoderm-specific genes that significantly increased in burst size than burst frequency whereas more mesoderm-specific genes changed in burst frequency (Fig. S8e,f).

Since histone acetylation has been suggested to influence transcription by modulating burst frequency ^68,69^, we sought to correlate tissue specific differences in burst kinetics to our genomic datasets. For the enhancer-paired DV genes with a significant kinetic change (*n* = 29), we found that the combined tissue-specific chromatin state at enhancers (*n* = 58) was a good predictor of changes in burst frequency between tissues (*R^2^* = 0.54), and correlated better with burst frequency changes than histone acetylation, CBP occupancy or chromatin accessibility individually (Fig. 6g and Fig. S8g). In contrast, the chromatin state at promoters was poor at predicting changes in burst frequency. Enhancer P-TEFb, BRD4/fs(1)h and eRNA transcription also correlated significantly with burst frequency, but were not as good predictors as the combined chromatin score (Fig. 6g). Differences in burst size between tissues could not be explained as well as burst frequency by the chromatin state, but a significant correlation of moderate strength was noted at enhancers (Fig. 6g and Fig. S8g, *R^2^* = 0.25). Interestingly, loading of CycT at promoters was among the best predictors of burst size, indicating that release from pausing may influence the burst size (Fig. 6g). We also explored if the enhancer-promoter distance influenced the modulation of burst size or frequency during activation. Interestingly, for a sub-class of DV enhancers located proximal (< 700 bp) to their target promoters (*n* = 22) (Fig. 6h), bursts involved a significantly stronger shift in size than frequency, whereas genes regulated by distal enhancers (*n* = 115) shifted in both burst size and frequency upon activation (Fig. 6i).

To further explore DV gene bursts, we plotted the kinetics for genes partitioned into classes based on whether they significantly changed in burst size (*n* = 8 genes, *n* = 19 enhancers), burst frequency (*n* = 16 genes, *n* = 33 enhancers) or both (*n* = 5 genes, *n* = 6 enhancers) in the cell cluster of expression compared to the inactive tissues (Fig. S8h-i). Burst frequency (k_on_) and burst size (k_syn_/k_off_) can reliably be inferred from scRNA-data ^68^, but how well the individual k_syn_ and k_off_ parameters can be estimated is more uncertain. We observed that increases in burst size appear to occur from lower off rates (*k*_off_) and not from increases in the rate of transcription (k_syn_) in the tissue of activity (Fig. S8h-i). Although there is uncertainty in these parameters, the data indicate that genes with increased burst size may remain in the ON state for a longer period of time when they are activated. The parameters of promoter mean occupancy (*k*_on_/(*k*_on_+*k*_off_)), which is the probability of a gene being in the ON state; switching correlation time (1/(*k*_on_+_*k*off_)), which is the correlation time of the switching process between ON and OFF states, and the mean transcript synthesis rate ((*k*_syn_/*k*_on_)/(*k*_on_+*k*_off_)); the rate of mRNA synthesis in the ON state ^70^, are also uncertain. Nevertheless, the data indicate that increases in either burst size or frequency result in similar transcript synthesis rates and that DV genes that change in both burst size and frequency achieve the highest rates of synthesis (Fig. S8h-i). Examining the correlations between each parameter and the chromatin state suggests that for genes changing in burst size, the enhancer chromatin state correlates well with burst size (*R^2^* = 0.48) and promoter mean occupancy (*R^2^* = 0.49) (Fig. S8j). This suggests that while the enhancer chromatin state primarily influences burst frequency, it can also modulate transcriptional bursts through other parameters in a context-dependent manner.

Taken together, our transcriptome-wide inference of transcriptional bursting dynamics during DV patterning show that DV genes have the capacity for high burst size, but a lower burst frequency than non-DV genes, and that individual genes vary in their dependencies on changes in size and frequency kinetics during bursts. Combining the burst data with our comprehensive genome-wide epigenomic data reveals that the enhancer chromatin state strongly modulates burst frequency, but has less, although still significant, influence on burst size. The high burst size of DV genes is encoded by core promoter motifs that mediate strong Pol II recruitment, as well as promoter-proximal pausing. Tissue-specific P-TEFb recruitment ensures that bursts are only triggered in specific cells. Exploring the burst strategies employed by DV genes reveals that despite varied dependencies of bursts for changes in frequency and size, similar rates of transcript synthesis are achieved in cells where the DV genes are active.

## Discussion

The establishment and maintenance of differential gene expression programs allows cells within multicellular organisms that contain genomes with identical DNA sequences to form distinct specialised tissues during embryogenesis. Yet, the interplay between chromatin state and transcription is not entirely understood. Here we have provided a comprehensive genome-wide assessment of chromatin state during *Drosophila* DV patterning, as measured by histone acetylation, chromatin accessibility and CBP occupancy, and directly compared it to zygotic transcription and Pol II activity status. The use of homogenous DV tissue mutants invariant in chromatin state and transcription, allowed us to dissect the interplay between the two. Consistent with data from mammals ^29^, we find that the chromatin state at promoters is largely similar across tissues and cell types, but that enhancers are marked by tissue-specific chromatin accessibility, histone acetylation and CBP occupancy.

This indicates that CBP fulfils distinct roles at enhancers and promoters. At enhancers, CBP is recruited and activated by dimerization induced by tissue-specific TFs to catalyze H3K27ac ^71^, which activates enhancers and stimulates target gene transcription. At promoters, CBP functions in the recruitment and establishment of a paused Pol II, possibly by interactions with the general transcription factor TFIIB ^33^. These results suggest that detection of CBP at the promoter is not simply a result of looping of the promoter to CBP-bound enhancers, as CBP can be enriched at the promoters of DV genes in a tissue in which it is absent at the enhancer. Further work is needed to elucidate the mechanisms underlying the deployment of distinct CBP activities at promoters and enhancers.

Inactivity in non-expressing tissues is sometimes mediated by active repression. Interestingly, we found that the chromatin state differs between two types of repressors, the short-range repressor Sna and the long-range repressor Dl. Sna-mediated repression involves a block of CBP-mediated H3K27ac, whereas repression by Dl is accompanied by Polycomb silencing and H3K27me3. Since PRC1 directly inhibits the HAT activity of CBP ^72^ and CBP-mediated acetylation of the +1 nucleosome stimulates Pol II release into elongation ^73^, PcGs may also target CBP to impede pause release of Pol II.

Interestingly, our results show that Pol II pauses at the promoters of DV genes in a tissue-invariant manner, irrespective of future transcription activation. Pol II promoter pausing has previously been shown to be an important regulatory step in the transcription of developmental genes and has been suggested to prime developmental genes for subsequent activation ^10^. Another function for Pol II pausing could be to promote synchronous gene activation across cells in a tissue ^12^, and to minimize expression variability between cells in a tissue ^24^. This property is largely defined by core promoter elements, with paused and synchronous genes often having Initiator (Inr), downstream promoter element (DPE) and pause button (PB) sequences, whereas TATA-containing genes show higher variability in expression and are less paused ^10,24^.

We find that pause release of Pol II is the critical regulatory checkpoint that dictates differential gene expression along the DV axis. Pausing is associated with the negative elongation factor (NELF) and DRB sensitivity inducing factor (DSIF, consisting of Spt4 and Spt5), and release of paused Pol II into productive elongation requires recruitment of the positive transcription elongation factor b (P-TEFb) kinase ^50^. P-TEFb phosphorylates NELF, DSIF and the C-terminal domain (CTD) of the Pol II largest subunit, allowing Pol II to escape from the pause site. Thus, control of P-TEFb recruitment and activity could be the key event in embryonic DV patterning. Indeed, we find that P-TEFb recruitment is spatially and temporally linked to DV gene activity. Interestingly, REL-family proteins such as NFκB regulate genes by targeting P-TEFb and transcription elongation in mammals ^74^, suggesting that the REL-protein Dl may also specify dorsoventral cell fates primarily by promoting pause release. In addition to transcription factors such as Dl, enhancer chromatin state could also influence pause release. We have previously shown that increased histone acetylation leads to release from pausing at a subset of genes ^73^, so the correlation we find between H3K27ac and tissue-specific gene activity may promote transcription elongation. It will be interesting to further investigate how signals from the enhancer can modulate the activity of P-TEFb.

Once a gene is turned on, transcription is not continuous, but occurs in bursts. We found that compared to other genes, DV genes have a low burst frequency but a high burst size. Thus, many transcripts are produced per burst. This may result from an enrichment of core promoter motifs in DV genes and high promoter-proximal pausing. However, pausing may also represent an alternative OFF state that is not captured by a two-state model of transcription ^75,76^. The majority of DV genes have a higher burst size in their tissue of expression compared to non-expressing cells. The burst size is determined by the initiation rate and the off rate. Unlike gap genes in *Drosophila* nc 13 embryos where the initiation rate is constant ^70^, our transcriptome-wide analysis showed that initiation rates vary between genes and between cells for individual genes. Still, the initiation rate (k_syn_) is less variable than the off rate (k_off_) for genes that increase their burst size in the tissue of expression. Burst size has been shown to increase in response to Notch signaling ^77,78^, primarily due to an increased burst duration. We cannot fully explain the increase in burst size in cells where DV genes are expressed, but we note that proximal enhancers and the presence of P-TEFb at the promoter may play a role, as well as the chromatin state at enhancers. The difference in chromatin state at enhancers between cells has an even larger impact on the burst frequency, consistent with previous findings ^68,69^, and with a role for enhancers in modulating burst frequency ^79^.

Surprisingly, genes that are believed to be regulated in the same fashion have different bursting kinetics. Both *twi* and *sna* are activated by the Dl transcription factor in the mesoderm, but whereas *twi* has a higher burst frequency in the mesoderm compared to neuroectoderm and dorsal ectoderm, *sna* expression is driven by a higher burst size. Further, both *dpp* and *zen* are directly repressed by Dl in neuroectoderm and mesoderm, but whereas burst frequency is increased for *dpp*, the *zen* burst size increases in dorsal ectoderm. Regulation of promoter occupancy (k_on_/(k_on_ + k_off_)), i.e. the proportion of time the promoter is active, has been suggested to establish the expression domains of the *Drosophila* gap genes ^70,80^, and two DV genes that respond to Dpp signaling ^81^. Consistent with this, we find that promoter occupancy is higher in cells where DV genes are expressed compared to other cell types both for genes that change their burst size and for those that change in frequency. However, we note that the kinetic parameters inferred in the framework of the two-state model may not be sufficient to fully explain the gene expression difference between cell types. Modulation of the window of time over which each cell transcribes the gene is a regulatory strategy that is independent of bursting, and important for *even-skipped* stripe 2 formation ^82^, which could also contribute to differential DV gene transcription.

Overall, these results augment our current understanding of the interplay between the formation of chromatin state and transcription (Fig. 6j). The data suggest that tissue-specific DV enhancers and promoters are initially primed by increased accessibility across nuclei prior to ZGA by the action of the maternally supplied pioneer factor Zelda ^61,83^. Increased accessibility at enhancers is accompanied by CBP recruitment and histone acetylation, priming the genes for future activation. This provides amenability for recruitment of Dl, with occupancy occurring differentially across the DV axis of the embryo according to its nuclear concentration and enhancer-specific differences in motif composition that affect binding affinity. Dl leads to the tissue-specific recruitment of other TFs and co-regulators, including CBP and BRD4, leading to the adoption of distinct enhancer chromatin states spatially within the embryo. Concomitantly, promoters become accessible across tissues, permitting the recruitment of CBP and Pol II by unidentified factors. Pol II initiates transcription and pauses before the Dl gradient has formed and remains paused in all tissues. Recruitment and activation of P-TEFb, likely mediated by Dl and distinct enhancer chromatin states, leads to tissue-specific pause release and differential gene expression. The frequency of transcriptional bursts (*k*_on_), is to a large part determined by the enhancer chromatin state, whereas the burst size (*k*_syn_/*k*_off_) may also depend on Pol II pausing and P-TEFb. We speculate that Pol II pausing confers a low off-rate, and that P-TEFb activity is regulated and important for a high synthesis rate, leading to a high burst size in the tissue of expression.

## Methods

### Drosophila stock maintenance

Mutant *Drosophila melanogaster* embryos composed entirely of presumptive dorsal ectoderm, neuroectoderm or mesoderm were obtained from the fly stocks *gd^7^/winscy hs-hid*, *Toll^rm9/rm10^*/*TM6 e Tb Sb* and *Toll^10B^/TM3 e Sb Ser/OR60*, respectively. One day old larvae laid by *gd^7^/winscy hs-hid* were heat shocked for 1.5 hr at 37°C for two consecutive days to eliminate *gd^7^* heterozygous animals and presumptive dorsal ectoderm mutant embryos collected from the remaining *gd^7^* homozygous flies. *Toll^rm9/rm10^* trans-heterozygous females were separated from the stock and from them presumptive neuroectoderm embryos collected. *Toll^10B^/TM3 e Sb* Ser and *Toll^10B^/OR60* heterozygotes that produce embryos composed of presumptive mesoderm were separated from the stock. Survival assays were performed to confirm embryonic lethality of *Toll* mutants. *yw; PCNA-eGFP*, a kind gift of Eric Wieschaus ^60^, and *w^1118^* lines served as controls (wild-type) for ChIP-qPCR and RT-qPCR experiments. Flies in which the *dorsocross* (*doc*) locus E1 enhancer had been deleted (doc enh del^Δ/Δ^) using a CRISPR (clustered regularly interspaced short palindromic repeats)-Cas9 mediated deletion strategy and an intermediary line carrying flippase recognition target (FRT) sites flanking the intact E1 enhancer (doc enh^+/+^) that served as a control, were kind gifts from Mounia Lagha ^8^. Flies carrying *UASp-CycT* and *Cdk9* transgenes were crossed with *w; alphaTub67C-GAL4∷VP16* (Bloomington line 7062) and used for maternal P-TEFb overexpression (OE) (see ‘Overexpression of P-TEFb in early embryos’ methods section).

Stocks were kept on potato mash-agar food and maintained at 25°C with a 12-hour light/dark cycle. Embryos were collected on apple juice plates supplemented with fresh yeast and aged at 25°C for specific time ranges dependent on the specific experiment which is detailed in the relevant methods section. Plates containing embryos collected for the first 2 h each day were discarded to avoid contamination by older embryos withheld by females. Collected embryos were dechorionated in diluted bleach, rinsed thoroughly in embryo wash buffer (PBS, 0.1% Triton X-100) and processed further in a manner dependent on the specific experiment which is detailed in the relevant methods sections.

### Overexpression of P-TEFb in early embryos

Coding sequences for the P-TEFb subunits CycT and Cdk9 were PCR amplified and cloned into the pUAS-K10.attB vector ^84^ by restriction digest and ligation (see Supplemental Table 9 for primer sequences) to produce the plasmids ‘*pUASp-CycT*’ and ‘*pUASp-Cdk9*’. CycT was cloned into pUAS-K10.attB using the *KpnI* and *XbaI* restriction sites whereas Cdk9 was cloned via *NotI* and *XbaI* sites. Plasmids were sequence verified, purified with the NucleoBond Xtra Midi kit (Machery-Nagel, Cat. 740410.50) and pUASp-CycT inserted into the attP2 landing site and pUASp-Cdk9 inserted into attP40 (FlyORF Injection Service). A double homozygous *UASp-Cdk9 ; UASp-CycT* stock was established and maternal overexpression achieved by crossing virgin females with *w; alphaTub67C-GAL4∷VP16* (Bloomington line 7062) males. The resulting *UASp-Cdk9*/*alphaTub67C-GAL4∷VP16 ; UASp-CycT/ +* females were collected, crossed with male siblings and used for embryo collection. For RNA *in situ* hybridization and ChIP-qPCR, P-TEFb OE and wild-type (*w^1118^*) embryos were collected for 2 h and aged a further 2 h (2-4 h AEL) and for RNA extraction and RT-qPCR embryos were collected for 1 h and aged a further 1.5 h (1.5-2.5 h AEL).

### RNA *in situ* hybridization

RNA in situ hybridization was performed on wild-type (w^1118^), gd^7^, Toll^rm9/rm10^and Toll^10B^ (2-4 h AEL) embryos using digoxigenin-labeled antisense RNA probes against dpp, ind, twi and shn. Probes against dpp, zen, ind, sog, twi and sna were used on wild-type (w^1118^) and P-TEFb OE (2-4 h AEL) embryos and a probe against psq was used on wild-type (w^1118^) (2-4 h AEL) embryos. RNA in situ hybridization was performed as previously described 85,86. Embryos were observed on a Leica DMLB 100T microscope and images taken on a Leica DMC2900 camera. We counted wild-type (wt) and P-TEFb OE (OE) embryos stained for dpp (wt n = 143, OE n = 206), *zen* (wt *n* = 119, OE *n* = 197), *ind* (wt *n* = 139, OE *n* = 178), *sog* (wt *n* = 155, OE *n* = 131), *twi* (wt *n* = 128, OE *n* = 86), and *sna* (wt *n* = 180, OE *n* = 112) manually for normal and ectopic signal and calculated the odds ratio alongside Fisher’s exact test to measure the significance of differences in the number of ectopically stained embryos between genotypes. Images of RNA *in situ* hybridization for *zen, ush, SoxN, Meltrin, twi, sna* and *htl* in wild-type embryos were obtained from the BDGP database ^87–89^.

### Precision run-on sequencing (PRO-seq)

PRO-seq was performed on *Toll* mutant embryos collected for 0.5 h and aged for a further 2.5 h (2.5-3 h AEL) or 4.5 h (4.5-5 h AEL) and both PRO-seq and qPRO-seq were performed on naïve *yw; PCNA-eGFP* embryos collected for 20 min, aged for 1 h (60-80 min AEL) and hand-sorted according to the nuclear cycle observed by the eGFP signal with older embryos discarded. Collected embryos were dechorionated in dilute bleach and rinsed thoroughly in embryo wash buffer (PBS, 0.1% Triton X-100) before being flash-frozen in liquid nitrogen and stored at −80°C.

PRO-seq and qPRO-seq were performed as previously described ^18,19^. Briefly, embryos were resuspended in cold nuclear extraction buffer A (10 mM Tris-HCl pH 7.5, 300 mM sucrose, 10 mM NaCl, 3 mM CaCl_2_, 2 mM MgCl_2_, 0.1% Triton X, 0.5 mM DTT, protease inhibitor cocktail (Roche) and 4 u/ml RNase inhibitor (SUPERaseIN, Ambion)), transferred to a dounce homogenizer and dounced with the loose pestle for 20 strokes. To remove large debris, the suspension was passed through mesh followed by douncing with a tight pestle for 10 strokes. Nuclei were pelleted at 700 *g* for 10 min at 4°C and washed twice in buffer A and once in buffer D (10 mM Tris-HCl pH 8, 25% glycerol, 5mM MgAc_2_, 0.1 mM EDTA, 5 mM DTT). For PRO-seq, isolated nuclei corresponding to approximately 10 million cells, and for qPRO-seq from 1 million cells, were resuspended in buffer D and stored at −80°C. Nuclear run-on assays were performed in biological duplicates exactly as previously described ^18,19,33^. PRO-seq and qPRO-seq libraries were sequenced (single-end 1 × 75 bp) on the Illumina NextSeq 550 platform at the BEA core facility, Karolinska Institutet, Stockholm.

### Assay for Transposase-Accessible Chromatin using sequencing (ATAC-seq)

ATAC-seq was performed on *Toll* mutant embryos collected for 0.5 h and aged accordingly to achieve three developmental time points: 2.5-3 h, 3.5-4 h and 4.5-5 h AEL. For each time point, 10 embryos per replicate presenting the correct morphology for the developmental stage sought were immediately hand-sorted. Hand-sorted embryos were dechorionated in dilute bleach, rinsed thoroughly in embryo wash buffer (PBS, 0.1% Triton X-100) and crude nuclear extracts isolated by homogenizing the embryos using a motor pestle in ATAC lysis buffer (10 mM Tris pH 7.4, 10 mM NaCl, 3 mM MgCl_2_ and 0.1% IGEPAL CA-630) and centrifugating at 700 *g* for 10 min. The nuclear pellet was resuspended in 22.5 μl of ATAC lysis buffer, 2.5 μl Tn5 (Tagment DNA Enzyme 1 (TDE1) (Illumina)) and 25 μl Tagment DNA Buffer (Illumina) and subjected to tagmentation at 37°C on a thermomixer at 1,000 rpm. Transposition was blocked by the addition of 1% SDS and DNA purified with Agencourt AMPure XP beads (Beckman Coulter, A63881) according to the manufacturer’s instructions using a 2:1 ratio of beads to sample. Libraries were prepared as previously described ^90^. Briefly, tagmented DNA was PCR amplified using 1x Phusion® High-Fidelity PCR Master Mix with GC Buffer (NEB) and 1.25 μM i5 and i7 PCR primers (Nextera® Index Kit (Illumina)) with the following PCR amplification conditions: 72°C for 5 min, followed by 10 cycles of 98°C for 10 seconds, 65°C for 1 min and 15 seconds, then 72°C for 1 min. Amplified libraries were purified with Agencourt AMPure XP beads with a 1.5:1 ratio of bead to sample volume. Libraries prepared from biological triplicates were sequenced paired-end (2 x 150 bp) on the Illumina NovaSeq platform at SciLifeLab, Stockholm.

### Chromatin immunoprecipitation sequencing (ChIP-seq) and ChIP-qPCR

ChIP-seq and ChIP-qPCR were performed on *Toll* mutant embryos collected for 2 h and aged for a further 2 h (2-4 h AEL) and ChIP-qPCR was also performed on *doc* enh del^Δ/Δ^ and *doc* enh^+/+^ (2-4 h AEL) control embryos and P-TEFb OE and wild-type (*w^1118^*) (2-4 h AEL) embryos. Formaldehyde crosslinking and chromatin preparation of embryos was performed as described previously ^91^. Briefly, dechorionated embryos were crosslinked in a mixture of 2 ml fixation buffer (PBS, 0.5% Triton X-100) and 6 ml heptane supplemented with 100 μl of 37% formaldehyde (Sigma-Aldrich, F8775) for 15 min at room temperature with rotation. Fixation was quenched by the addition of PBS supplemented with 125 mM glycine and crosslinked embryos were washed 3 times in wash buffer (PBS, 0.5% Triton X-100), snap frozen in liquid nitrogen and stored at −80°C. For chromatin preparation, embryos were homogenized in a glass dounce homogenizer by 20 strokes with a tight pestle in A1 buffer (15 mM HEPES pH 7.6, 15 mM NaCl, 60 mM KCl, 4 mM MgCl_2_, 0.5 mM DTT, 0.5% Triton X-100 and protease inhibitor tablets (Roche)), centrifuged at 3500 *g* for 5 min at 4°C and the supernatant discarded. The remaining nuclear pellet was resuspended in 200 μl of sonication buffer (15 mM HEPES pH 7.6, 140 mM NaCl, 1 mM EDTA pH 8.0, 0.5 mM EGTA pH 8.0, 0.1% sodium deoxycholate, 1% Triton X-100 and protease inhibitor tablets (Roche)) supplemented with 0.5% SDS and 0.2% n-lauroylsarcosine and sonicated using a Bioruptor (Diagenode) with high power settings to obtain an average fragment size distribution of 200-500 bp, Sonicated chromatin was centrifuged at 13,000 rpm for 10 min at 4°C and diluted 5-fold in sonication buffer to reduce the concentration of detergents.

For chromatin from *Toll* mutant embryos, immunoprecipitations (IPs) were performed with 10 μg rabbit anti-CBP (homemade, ^16^), 2 μg rabbit anti-CycT and anti-Cdk9 (both kind gifts of Kazuko Hanyu-Nakamura ^92^), and with 2 μg rabbit anti-H3K27ac (Abcam, ab4729) (also described in ^7^) and 5 μg mouse anti-H3K27me3 (Abcam, ab6002) on chromatin from the *Toll^rm9/rm10^* mutant. H3K27ac ChIP-seq data from *gd^7^* and *Toll^10B^* mutants used in this study were generated by ^14^. IPs on chromatin from *doc* enh del^Δ/Δ^ and *doc* enh^+/+^ control embryos were performed with CycT, rabbit anti-BRD4/fs(1)h (long isoform) (a gift of Renato Paro, kindly provided by Nicola Iovino) ^93^), CBP, H3K27ac, H3K27me3 and rabbit anti-H3 (Abcam, ab1791) antibodies. Chromatin from P-TEFb OE and wild-type (*w^1118^*) embryos was immunoprecipitated with anti-CycT. Chromatin was incubated with antibodies overnight at 4°C and equal amounts of Protein A and G Dynabeads (Invitrogen), pre-blocked with BSA (1 mg/ml), were incubated with the samples for 4 h at 4°C. Chromatin corresponding to 10% of the amount in each IP was withdrawn to serve as an input for qPCR. Samples were subjected to 10 min washes with Wash A (10 mM Tris-HCl pH 8.0, 1 mM EDTA pH 8.0, 140 mM NaCl, 0.1% SDS, 0.1% Sodium Deoxycholate and 1% Triton X-100), Wash B (Wash A adjusted to 500 mM NaCl), Wash C (10 mM Tris-HCl pH 8.0, 1 mM EDTA pH 8.0, 250 mM LiCl, 0.5% Sodium Deoxycholate and 0.5% IGEPAL CA-630) and Tris-EDTA (TE) buffer. Beads were resuspended in 100 μl TE and treated with RNase A (20 μg/ml) at 55°C for 30 min before SDS (to 0.75%) and Tris-HCl (to 50mM) were added and crosslinks reversed at 65°C for overnight. Eluted ChIP DNA was treated with Proteinase K at 55°C for 2 h and purified with the ChIP DNA Clean & Concentrator kit (ZymoResearch, D5205).

ChIP-sequencing was performed using 2-5 ng of ChIP DNA from *Toll* mutant IPs with CBP, H3K27ac and H3K27me3 antibodies (only *Toll^rm9/rm10^* for H3K27ac and H3K27me3 ChIP DNA) in biological duplicates. Libraries were prepared with the NEBNext® Ultra II DNA Library Prep Kit for Illumina (NEB, E7645L) and single-end (1 x 75 bp) sequenced on the Illumina NextSeq 550 platform at the BEA core facility, Karolinska Institutet, Stockholm.

ChIP DNA from IPs and inputs on *Toll* mutant chromatin (CycT), *doc* enh del^Δ/Δ^ and *doc* enh^+/+^ control chromatin (CycT, BRD4/fs(1)h, CBP, H3K27ac, H3K27me3 and H3) and P-TEFb OE and wild-type (*w^1118^*) chromatin (CycT) were analyzed by qPCR on a CFX96 Real-Time System (BioRad). qPCR reactions were carried out using 2 μl of ChIP DNA as a template with 300 nM primers and 5X HOT FIREPol® EvaGreen® qPCR Mix Plus (Solis BioDyne) in duplicate. All primers used in this study are listed in Supplemental Table S9 The percentage of input precipitated for each target was determined by comparing the average Cq to that of the input and the level of enrichment normalized to the signal at intergenic loci devoid of chromatin factors and histone modifications. For H3K27ac and H3K27me3, enrichment was further normalized to the occupancy of H3. Due to unexpected variations in the intergenic control signal between CycT IPs on P-TEFb OE and wild-type (*w^1118^*) chromatin, data were presented as percent (%) input precipitated.

### CUT&Tag

CUT&Tag was performed on *Toll* mutant embryos collected for 2 h and aged for a further 2 h (2-4 h AEL) and *yw; PCNA-eGFP* embryos collected for 20 min and aged for 1 h (60-80 min AEL, nc 7-9), 30 min and aged for 1.5 h (1.5-2 h AEL, nc 11-13) and 2 h and aged for 2 h (2-h AEL, nc 14). Hand-sorting was performed with the nuclear cycle observable by the eGFP signal with older embryos discarded. CUT&Tag was performed essentially as described by ^94^. Collected embryos were dechorionated, rinsed in embryo wash buffer (PBS, 0.1% Triton X-100) and crude nuclear extracts prepared using a glass douncer and loose pestle in Nuclear Extraction buffer (20 mM HEPES pH 7.9, 10 mM KCl, 0.5 mM spermidine, 0.1% Triton X-100, 20% glycerol with protease inhibitor cocktail (Roche)) ^95^ and centrifuged at 700 *g* for 10 min at 4°C. The nuclear pellets were resuspended in Nuclear Extraction buffer. Nuclei corresponding to 50 embryos per reaction (2-4 h AEL), 100 embryos (1.5-2 h AEL) or 200 embryos (60-80 min AEL) were incubated with 30 μl of BioMag®Plus Concanavalin A beads (Polysciences) (prepared in Binding buffer (20 mM HEPES pH 7.5, 10 mM KCl, 1mM CaCl_2_, and 1 mM MnCl_2_)) on a nutator for 10 min at 4°C. Nuclei-bead complexes were resuspended in 100 μl Antibody buffer (20 mM HEPES pH 7.5, 150 mM NaCl, 0.5 mM spermidine, 0.05% digitonin, 2 mM EDTA pH 8.0 and 0.1% BSA supplemented with protease inhibitor cocktail (Roche)). For *Toll* mutant CUT&Tag, 1 μl of rabbit anti-BRD4/fs(1)h, rabbit anti-CycT, rabbit anti-Cdk9, rabbit anti-Rpb3 (a kind gift of John Lis), guinea pig anti-Dorsal (a kind gift of Christos Samakovlis) and rabbit anti-RNA Polymerase II CTD repeat YSPTSPS (phosphor S5) (5SerP) (Abcam, ab5131) overnight at 4°C. For *yw; PCNA-eGFP* CUT&Tag, 1 μl of rabbit anti-BRD4/fs(1)h, rabbit anti-Cdk9 and rabbit anti-CBP. Following overnight incubation, tagmentation was performed using pA-Tn5 (Protein Science Facility, KI, Stockholm). Tagmented DNA was PCR amplified using custom i5 and i7 PCR primers and Phusion® High-Fidelity PCR Master Mix with GC Buffer (NEB). PCR conditions were: 72°C for 5 min, 98°C for 30 s, followed by thermocycling (98°C for 10 s and 63°C for 10 s) for 13 cycles and final extension at 72°C for 1 min. Amplified libraries were purified using Agencourt AMPure XP beads (Beckman Coulter) (1.1:1 bead to sample volume ratio). Libraries were paired-end (2 x 37 bp) sequenced on an Illumina NextSeq 550 platform at the BEA core facility, Karolinska Institutet, Stockholm. The low read counts obtained from sequencing for the CUT&Tag samples *gd^7^* 2to4h CUT&Tag CycT Replicate 2, *gd^7^* 2to4h CUT&Tag BRD4 Replicate 2 and *Toll^10B^* 2to4h CUT&Tag Dl Replicate 2 indicated these reactions had failed so they were excluded from the subsequent analysis.

### RNA extraction and RT-qPCR

Total RNA was extracted from *doc* enh del^Δ/Δ^ and *doc* enh^+/+^ (2-4 h AEL) and P-TEFb OE and wild-type (*w^1118^*) (1.5-2.5 h AEL) embryos. Dechorionated embryos were homogenized in cold PBS with a plastic pestle and RNA extracted using TRIzol LS (Invitrogen). Total RNA was purified and concentrated using the RNeasy MinElute Cleanup kit (Qiagen) according to the manufacturer’s instructions. Purified RNA (1.5 μg) was treated with DNase I (Sigma-Aldrich) to eliminate contaminating genomic DNA and converted to cDNA with the High-Capacity RNA-to-cDNA kit (ThermoFisher Scientific) according to the manufacturer’s instructions. RT-qPCR was performed on a CFX96 Real-Time System (Biorad) using 2 μl of cDNA as template with 300 nM primers and 5X HOT FIREPol® EvaGreen® qPCR Mix Plus (Solis BioDyne) in duplicate. The delta-delta Ct method was used to quantify mRNA levels relative to RpL32 (RNA from *doc* enh del^Δ/Δ^ and *doc* enh^+/+^ embryos) and 28S rRNA (RNA from P-TEFb OE and wild-type (*w^1118^*)). All primers used in this study are listed in Supplemental Table S9.

### PRO-seq data analysis

PRO-seq and qPRO-seq reads were mapped to the *Drosophila melanogaster* (dm6) genome assembly using Bowtie2 (v.2.3.5) with the default program parameters ^96^. Library mapping statistics are listed in Supplemental Table S10. Strand separated RPKM normalized (bigwig) coverage tracks from individual replicates were generated using the deepTools (v.3.5.1) package ‘bamCoverage’ using the default parameters (binSize = 2 (bases), normalizeUsing = RPKM) ^97^. Strand separated files of the mean RPKM signal from both replicates were produced by first merging the read alignment files produced by Bowtie2 from each replicate using the SAMtools package ‘samtools merge’ and then bigwig files produced by ‘bamCoverage’ (deepTools). To allow for simultaneous genome browser visualization of the signal from the pause site and gene body at genes of interest, the bin size was extended to 10 bp when producing the merged bigwig files. Read counts per gene (CDS) were extracted with featureCount ^98^.

### ChIP-seq, ATAC-seq and CUT&Tag data analysis

ChIP-seq, ATAC-seq and CUT&Tag reads were mapped to the *Drosophila melanogaster* (dm6) genome assembly using Bowtie2 (v.2.3.5) with the default program parameters ^96^. Library mapping statistics are listed in Supplemental Table S10. RPKM normalized (bigwig) coverage tracks from individual replicates were generated using the deepTools (v.3.5.1) package ‘bamCoverage’ using the default parameters (binSize = 1 (bases), normalizeUsing = RPKM) ^97^. The mean RPKM signal from both replicates were produced by first merging the read alignment files produced by Bowtie2 from each replicate using the SAMtools package ‘samtools merge’ and then bigwig files produced by ‘bamCoverage’ (deepTools). Peaks were called for *Toll* mutant ATAC-seq and CBP and H3K27ac ^7,13,14^ ChIP-seq using the Genrich peak caller (version 0.6) (https://github.com/jsh58/Genrich#contact) with the default program parameters. Read counts per gene (CDS), promoter and enhancer (all peaks called for CBP not overlapping the TSS of genes) were extracted with featureCount ^98^.

### Analysis of previously published datasets

In addition to the datasets generated in this study we reanalyzed the following published datasets: ChIP-seq for H3K27ac and H3K27me3 from *gd^7^* and *Toll^10B^*, Zen and Mad from *gd7* embryos, Twi from *Toll^10B^*, Sna, Pc and GAF from wild-type (Oregon-R) (2-4 h AEL) embryos (GEO: GSE68983) ^14 13^; ChIP-seq for H3K27ac from *Toll^rm9/rm10^* (2-4 h AEL) embryos (ArrayExpress: E-MTAB-9303) and scRNA-seq from wild-type (*PCNA-eGFP* and *w^1118^*) and *Toll* mutant (2.5-3.5 h AEL) embryos (ArrayExpress: E-MTAB-9304) ^7^; ChIP-nexus for Dl from wild-type (Oregon-R) (2-4 h AEL) embryos (GEO: GSE55306) ^41^; ChIP-seq for Zld from nc 8, nc 13 and nc 14 wild-type embryos (GEO: GSE30757) (Harrison et al., 2011); ChIP-seq for Zld from wild-type (2-3 h AEL) embryos (GEO: GSE65441) ^61^; ChIP-seq for Opa from wild-type (ZH-86Fb) nc 14 (4 h AEL) and ATAC-seq from nc 14 wild-type (ZH-86Fb), Zld maternal RNAi and opa maternal RNAi embryos (GEO: GSE140722) ^63^; CLAMP ChIP-seq for CLAMP from wild-type (MTD-Gal4, Bloomington line 31777) (2-4 h AEL) embryos (GEO: GSE152598) and ATAC-seq from wild-type (MTD-Gal4, Bloomington line 31777) and CLAMP maternal RNAi (2-4 h AEL) embryos (GEO: GSE152596) ^64^; ATAC-seq from control (His2AV-RFP; sfGFP-GAF) and GAF^deGradFP^ (His2Av-RFP/nos-degradFP; sfGFP-GAF) (2-2.5 h AEL) embryos (GEO: GSE152771) ^65^; ChIP-seq for H3K27ac, H3K18ac, H4K8ac, H3K9ac, H3K4me1 and H3K4me3 ChIP-seq from wild-type (Oregon-R) (nc 8, nc 12 and nc 14 (early and late)) embryos (GEO: GSE58935) ^66^; and ATAC-seq from wild-type embryos nc 11-13 (GEO: GSE83851) ^60^.

Reads for the publicly available data were mapped to the *Drosophila melanogaster* (dm6) genome assembly using Bowtie2 (v.2.3.5) with the default program parameters ^96^. RPKM normalized (bigwig) coverage tracks from individual replicates were generated using the deepTools (v.3.5.1) package ‘bamCoverage’ using the default parameters (binSize = 1 (bases), normalizeUsing = RPKM) ^97^. For ATAC-seq data from wild-type embryos ^60^, bigwig files of the mean signal for replicates and the mean signal across each nuclear cycle were produced using the deepTools (v.3.5.1) package ‘bigwigCompare’ using the default parameters. For ChIP-seq data for various histone modifications ^66^ and ATAC-seq data for from various pioneer factor perturbations ^63–65^ we used processed data sets generated in the original publications.

### Quality control for PRO-seq, ATAC-seq, ChIP-seq and CUT&Tag

For *Toll* mutant PRO-seq, ATAC-seq, ChIP-seq and CUT&Tag experiments, Principal Component Analysis (PCA) was done on the normalized read counts for all genes, promoters and enhancers (peaks called for CBP not overlapping the TSS of genes) as a quality control (QC) step to make sure that most of the variation in the data could be explained by the difference in genotype between the mutants. Based on the scores of the PCA, a subset of the principal components (PCs) were identified that separated the *Toll* mutant samples in the PC space.

### Identification of tissue-specific regions linked to DV regulated genes

Based on the PCs identified in the QC, three latent linear vectors, one for each *Toll* mutant, were created. For each *Toll* mutant, the vectors pass through origo and the mean value of the *Toll* mutant samples PC scores with the positive direction towards the mean value of the *Toll* mutant. For each region (genes, promoters and enhancers), three latent vector scores, one for each *Toll* mutant, were calculated. Each score is the position on the *Toll* mutant latent vector that is the closest to the regions PCA loading. For each *Toll* mutant the scores were then normalized, by removing the mean and dividing by the standard deviation of all the regions. For genes, the regions represent all expressed genes, for promoter it represents the regions around the expressed genes’ TSS and for enhancers it represents the peak regions identified from the CBP ChIP-seq peak calling.

### Identification of differentially expressed DV regulated genes and pausing analysis

Genes with less than 10 reads mapping to the gene body were removed from the analysis. Count data for the remaining genes were normalized using DEseq2 ^99^. Log_2_ normalized gene levels were used for quality control and latent vector scores as described above. To test the validity of using a latent vector approach and select a cutoff, ROC analysis was done. The positive set, for the ROC analysis, of previously known DV regulated genes identified from microarray data and validated by one other method were obtained from ^100^. After ROC analysis DV genes were selected with a latent vector score above 3. A list of AP regulated genes (*n* = 31) used as a comparative data set to the DV genes were obtained from Saunders, et al. ^101^. Genes zygotically expressed at nc 7-9 (*n* = 20), nc 9-10 (*n* = 63), syncytial blastoderm (*n* = 946) and cellular blastoderm (*n* = 3540) stages of early *Drosophila* embryogenesis were obtained from Kwasnieski, et al. ^22^.

To examine Pol II promoter proximal pausing, the gene body read count (GBC, using the CDS counts) and promoter count (PC, from 50 bp upstream of the TSS to 100 bp downstream of the TSS) were determined for each annotated transcript. The pausing index (PI), which is a ratio describing the magnitude of pausing, was calculated by dividing the PC by the sum of the PC and GBC. For genes with multiple isoforms, the transcript with the highest average GBC divided by the length of the CDS was selected. Statistical analysis for comparisons between the gene expression level and PI of the same gene class between different *Toll* mutants were performed with the Wilcoxon signed-rank test, whereas comparisons between different gene classes used the Wilcoxon rank-sum test.

### Identification of tissue-specific enhancers linked to DV regulated genes

To identify tissue specific enhancers regions, identified by CBP ChIP-seq peak calling, were analysed for enrichment of reads from CBP ChIP-seq, H3K27ac ChIP-seq and ATAC-seq *Toll* mutant experiments described above. For each approach regions with less than 10 reads were removed from further analysis and the remaining regions were normalised with DEseq2 rlog and used for PCA analysis. Number of PC for each approach was manually selected, PC 1 to 3 for ATACseq and PC 1 and 2 for CBP and H2K27ac and used for latent vector score calculations as described above. Combined tissue-specific enhancer and promoter latent vector scores were calculated by summing the CBP, H3K27ac and the ATAC-seq enhancer or promoter latent vector scores for each *Toll* mutant. Only the enhancer regions among the top 5% were considered potential enhancers. When assigning putative enhancers to DV regulated genes, a requirement was that they resided in the same topologically associated domain (TAD) (domain boundaries were from Hi-C data in 3-4 h AEL embryos ^7,42^.

To validate the functional activity of the identified tissue-specific DV enhancers, we lifted annotation terms (*n* = 31) associated with the *in vivo* activity of 7793 enhancer reporter lines driven by non-coding genomic fragments in stage 4 to 6 *Drosophila* embryos ^32^. We then measured the enrichment of annotation terms for reporter lines driven by fragments overlapping DV enhancers and compared to those overlapping all other annotated CBP peaks. Only terms with *P*-values < 0.005 (Fisher’s exact test) in at least one of the *Toll* mutant enhancers were kept.

To assess the quality of the enhancer identification strategy we performed receiver operating characteristic (ROC) curve analysis with all non-assigned enhancer regions as the negative set and the assigned tissue specific enhancers that overlapping non-coding genomic fragments identified to have DV-regulated activity in an enhancer-reporter assay ^32^ as the positive set.

### Promoter and enhancer motif analysis

We scanned DV promoters for putative core promoter elements from the CORE database and compared the proportion of promoters with motifs between DV promoters and all promoters in the database ^26^. To *de novo* identify promoter motifs we scanned DV regulated promoters for ungapped enriched motifs using the Multiple EM for Motif Elicitation (MEME) tool from the MEME suite (https://meme-suite.org/meme). Long enriched motifs were identified with a threshold of 31 nt (-minw 31). For the other motifs we required that the length to be ≤ 30 nt (-maxw 30). Motifs with an e-value less than 0.005 were kept for further analysis. The motifs were compared, using TOMTOM in the MEME-suite, against the motifs in the JASPAR Insects CORE redundant TF motifs database (version 2020). All of the *de novo* identified motifs with *P*-values less than 0.005 were renamed based on matches to known motifs from the JASPAR database. Motifs that fitted the Inr and the DPE were assigned as Inr or Inr and DPE. The MEME suite tool FIMO was used to search DV, AP and all other promoters for occurrences of the identified motifs. Log_2_ odds ratios were measured for the different motifs. See Table S3 for the *de novo* identified motifs at DV promoters.

The MEME tool was also used to identify motifs enriched at the tissue-specific DV enhancers. For each enhancer class, we scanned for motifs between 5-mer and 12-mer in width, occurring with any number of repetitions within each sequence. We then compared the *de novo* identified motifs to known *Drosophila* motifs across the combined *Drosophila* databases from the MEME suite using TOMTOM with the default parameters (Table S5). Enriched known motifs, from the JASPAR Insects CORE redundant TF motifs database (version 2020), were identified using the function ‘motifEnrichment’ from the PWMEnrich 4.26.0 R package (https://bioc.ism.ac.jp/packages/3.11/bioc/html/PWMEnrich.html). Five hundred randomly selected CBP peaks were used as a background distribution. All motifs with a raw enrichment score of > 1.5 and a *P*-value < 0.05 in at least one of the enhancer classes were kept as enriched motifs (Table S6).

### Examining DV enhancer and promoter overlaps with peaks called from publicly available data

We performed MACS2 peak calling ^102,103^ with the default program parameters on the alignments produced with Bowtie2 from the published ChIP-nexus for Dorsal (Dl) ^41^ and ChIP-seq for Zen, Mad, Twi and Sna ^14 13^. From the MACS2 peak calling results, we selected the 500 (Dl and Sna) or 1000 (Zen, Mad and Twi) strongest sites to ensure only high-confidence peaks would be used in the analysis. BEDTools intersect ^104^ was used to identify DV enhancers and promoters that overlapped the called peaks. For measuring overlaps, promoter regions were defined as (TSS ± 750 bp). DV genes identified from *Toll* mutant PRO-seq expressed specifically in the dorsal ectoderm with a Dl peak within ± 15 kb of the TSS were selected as Dl-repressed genes (*n* = 6). Sna repression targets (*n* = 13) were identified as the genes expressed in the neuroectoderm with Sna and Dl co-bound site within ± 15 kb of the TSS. Overlaps were also made using a list of genomic coordinates for known *Drosophila* Polycomb Response Elements (PREs) compiled by ^58^ and lifted to the dm6 *Drosophila* reference genome using the UCSC LiftOver tool. To preserve the original spike-in normalizations used for the ChIP-seq data from histone marks across early nuclear cycles ^66^, we used the coordinates for peaks called in the original paper using the dm3 *Drosophila* reference genome. We lifted peaks for DV and non-DV enhancers and promoters from dm6 to dm3 using the UCSC LiftOver tool. For Zld ChIP-seq across early nuclear cycles ^62^ we also used the peaks called in the original paper from the dm3 reference genome.

### Measuring changes in chromatin accessibility at DV enhancers and promoters after pioneer factor perturbation

To quantify changes in ATAC-seq signal at DV enhancers and promoters from publicly available data for pioneer factor perturbations ^63–65^, the signal at DV and shuffled enhancers and promoters (TSS ± 500 bp) (obtained using BEDTools ‘shuffle’^104^) was counted using the deepTools ‘BigWigSummary’ tool ^97^ and the log_2_ fold change (perturbation/control) in signal measured. Boxplots were produced in R using the ggplot2 package and significant differences in the change in accessibility between DV and shuffled enhancers and promoters was measured with the Wilcoxon rank-sum test.

### Uniform Manifold Approximation and Projection (UMAP) clustering of scRNA-seq data

We selected the cells in scRNA-seq from wild-type (*PCNA-eGFP*) embryos that had been originally assigned to DV-relevant clusters (ectoderm1, ectoderm2, ectoderm3, neural1, neural2, mesoderm1 and mesoderm2, *n* = 2787) based on clustering using the shared nearest neighbor (SNN) approach from the Seurat package (version 4.1.0) ^7,105,106^. From the scRNA-seq data in the selected cells, a new principal component space was constructed using the 195 PRO-seq identified DV genes as features to separate the cells. SNN clustering was performed on the first 10 PCs with a clustering resolution of 0.3. Identified clusters were annotated based on the expression levels of PRO-seq identified DV genes. Based on the expression of DV marker genes the derived clusters were named Dorsal ectoderm (*dpp, Doc1* and *ush* marker genes, *n* = 1396), early (*ind*, *sog* and *brk* marker genes, *n* = 1392) and late (older neural cells, *scrt*, *ase* and *nerfin-1* marker genes, *n* = 2367) Neuroectoderm, early (*twi and sna* marker genes, *n* = 2333) and late (older mesoderm or myoblasts, *Mef2*, *meso18E*, *sns* and *sing* marker genes, *n* = 1448) Mesoderm and a common cluster of cells (*n* = 851) that could not be separated according to the expression of the DV genes (Table S7). Average expression levels within each cluster were obtained for 160 of the 195 DV regulated genes, 26 of the 31 AP regulated genes and 1819 non-DV genes (Table S7). Uniform Manifold Approximation and Projections (UMAPs) were constructed using the default settings to visualize the scRNA-seq data.

### Inference of transcriptional bursting kinetics from scRNA-seq data

To infer transcriptional bursting kinetics, scRNA-seq UMI count matrices from the two wild-type samples (PCNA:eGFP and *w^1118^*) were first subsetted per cluster. For the dorsal ectoderm, neuroectoderm (early) and mesoderm (early) clusters, maximum-likelihood kinetics inference was attempted for all detected genes according to the model implemented by Larsson, et al. ^68^. Additionally, pseudorandom bootstraps of the input data before maximum-likelihood inference in 100 iterations were performed. Through the bootstrapped inference, empirical confidence intervals could be derived. Next, we filtered away low-power inferences outside of the parameter space by sorting the values into two distributions based on a mixture of two normal distribution curves using the normalMixEM tool from the mixtools package in R (version 1.2.0) (https://cran.r-project.org/web/packages/mixtools/vignettes/mixtools.pdf) and the values in the higher distribution were kept. Genes with noisy confidence inferences (i.e. a broad confidence interval (CI)) were removed (For *k*_on_: log_10_(CI *k*_on_) < 1.3 + 0.8 log_10_(*k*_on_) and for *k*_bs_: log_10_(CI *k*_bs_) < 1.0 + 0.8 log_10_(*k*_bs_). Kinetics were obtained for 2232 genes in all three clusters and 1519 genes in two of the three clusters. Genes where the CI for two clusters did not overlap were considered to be significantly different. Pearson correlations of the bursting kinetics for the DV clusters between the two wild-type samples were measured to control for reproducibility. DV genes were separated into kinetic classes based on whether they significantly changed in burst frequency (*n* = 16), burst size (*n* = 25), both burst size and burst frequency (*n* = 6) or did not change significantly (*n* = 83).

## Data and code availability

The datasets generated during this study are available at Gene Expression Omnibus with the Accession Number GEO: GSE211220.

## Acknowledgements

We thank Mounia Lagha, Christos Samakovlis, Akira Nakamura, John Lis, Renato Paro, Eric Wieschaus and Nicola Iovino for sharing reagents, Juanma Vaquerizas and Mounia Lagha for comments on the manuscript, the core facility for Bioinformatics and Expression Analysis (BEA) at Novum (supported by the board of research at the Karolinska Institute and the research committee at the Karolinska hospital) for help with sequencing, and NBIS/SciLifeLab for bioinformatics long-term support (WABI). Data handling was enabled by resources provided by the Swedish National Infrastructure for Computing (SNIC), partially funded by the Swedish Research Council through grant agreement no. 2018-05973. This work was supported by grants from the Swedish Research Council (Vetenskapsrådet) and the Swedish Cancer Society (Cancerfonden) to M.M.

## Author contributions

G.H. and R.V. performed experiments including PRO-seq, ATAC-seq, ChIP-seq as well as CUT&Tag with help from A.P. P-TEFb over-expression was performed by S.P. Bioinformatic analysis: J.R. with help from G.H. and R.V. Transcriptional kinetics: J.R., C.Z. and R.S. Conceptualization: M.M., R.V. and G.H. Writing: M.M. and G.H. with input from all authors.

---

**Figure S1. PRO-seq identifies DV regulated genes with promoter-proximal paused Pol II that persists across tissue types and developmental stages. a)** Fold change (log_2_ TPM +1) of PRO-seq gene body read counts (GBC) (defined as the coding region of genes) in *Toll* mutants for DV genes grouped by the tissue of expression and non-DV genes. P-values denoting significant differences for DV gene groups between *Toll* mutants are from the Wilcoxon signed-rank test. **b)** Counts for the differentially expressed genes identified by PRO-seq, grouped by the tissue of expression. **c)** The overlap (%) PRO-seq identified DV regulated genes with DV genes previously identified by whole genome microarray ^20^. **d)** Images of whole mount *in situ* hybridization in wild-type embryos (2-4 h AEL) with probes for mRNAs of representative DV regulated genes identified by PRO-seq. Images of *zen, ush, SoxN, Meltrin, twi, sna* and *htl* were obtained from the BDGP database ^87–89^. **e)** Genome browser shots of stranded PRO-seq signal (RPKM x10^3^) at *Wnt2*, *wnd* and *meso18E*. Promoters are shaded gray. **f)** Pausing index (PI) of DV regulated genes compared to anterior-posterior (AP) ^101^ and non-DV genes in *Toll* mutant PRO-seq. **g)** Comparisons of the transcription level (GBC (log_2_ TPM+1)) of zygotic genes expressed at different embryonic developmental stages ^22^ and DV genes between naïve wild-type embryos in qPRO- and PRO-seq. **h)** Metagene plots of naïve qPRO- and PRO-seq signal at developmentally staged and DV genes. **i)** Genome browser shots of stranded qPRO- and PRO-seq signal (RPKM x10^3^) from naïve wild-type embryos at representative nc 7-9 expressed genes. **j)** Comparisons of the PI of the gene classes from **g** between naïve wild-type embryos in qPRO- and PRO-seq. **k)** Representation (%) of core promoter elements from the CORE database ^26^ at the promoters of DV regulated (*n* = 195) and all (*n* = 13,965) genes. **l)** Venn diagrams of the overlap of DV regulated genes with Inr, DPE and Bridge motif. **m)** Representation (%) and odds ratio (log_2_) for *de novo* identified motifs at the promoters of DV, AP and other genes. **n)** De novo identified motif densities for Inr, Inr-DPE, Mad-Brk, GAGA and TATA at the promoters (from 100 bp downstream to 50 bp upstream of the TSS) of DV, AP and other genes. **o)** Metagene plots of *Toll* mutant CUT&Tag (2-4 h AEL) at DV regulated genes performed with antibodies against Pol II (Rpb3) and serine 5 phosphorylated (5SerP) Pol II. **p)** PRO-seq promoter counts (PC) (log_2_ TPM+1), GBC and PI for representative DV genes from 2.5-3 h and 4.5-5 h AEL *Toll* mutants.

**Figure S2. Characterization of tissue-specific DV enhancers identified by epigenomic profiling of chromatin state. a)** Counts for the tissue-specific DV enhancers and **(b)** corresponding promoters identified partitioned by the tissue of activity of target genes. **c)** Enhancer-linked DV genes binned by the number of paired enhancers. **d)** Distribution of DV enhancer density in relation to genomic distance from the TSS of target genes. Enhancers with a distance ≤ 700 bp from the TSS were classified as proximal and above this threshold defined as distal. **e)** Metagene profiles of CBP, ATAC-seq and H3K27ac enrichment at DV enhancers (± 5 kb of CBP peak). **f)** Violin plots of the genomic length distributions (kb) of peaks called for CBP, ATAC-seq and H3K27ac overlapping DV enhancers. **g)** Enriched annotations associated with the enhancer reporter activities of non-coding genomic fragments (Vienna Tiles, VT) that overlap DV enhancers partitioned by the tissue of activity^32^. **h)** Images of whole-mount *in situ* hybridization of LacZ reporter activity driven by representative VT enhancer fragments that overlap identified DV enhancers. **i)** Metagene profiles and heatmaps (± 5 kb of CBP peak) of *Toll* mutant H3K27ac and CBP ChIP-seq signal (RPKM) at dorsal ectoderm, neuroectoderm and mesoderm enhancers. **j)** Receiver operating characteristic (ROC) curves for ATAC-seq, CBP and H3K27ac individually and combined at DV enhancers and promoters**. k)** Metagene profiles and heatmaps (± 5 kb of TSS) of *Toll* mutant CBP and ATAC-seq signal (RPKM). **l)** Genome browser closeups of *Toll* mutant PRO-seq (3 h AEL) signal at the *twi* promoter and enhancer.

**Figure S3. Detection of tissue-specific transcription factors at DV enhancers and examination of genome organization. a)** Significantly enriched motifs from the JASPAR Insects CORE redundant TF motif database at DV enhancers, separated by the tissue of activity. **b)** Cumulative enrichment scores of the motifs from **a** at DV enhancers. **c)** Overlap (%) of peaks called for Dl, zen, Mad and twi with all DV enhancers and enhancers partitioned by the tissue of target gene activity. **d)** Normalized Hi-C contact probabilities (5-kb resolution) for representative DV regulated genes (*dpp*, *ind* and *twi*) in *Toll* mutant (2-3 h AEL) embryos from Ing-Simmons, et al. ^7^.

**Figure S4. Tissue-specific P-TEFb and BRD4/fs(1)h recruitment to DV genes. a)** Metagene profiles and heatmaps (± 5 kb of TSS) of *Toll* mutant CUT&Tag (2-4 h AEL) at DV regulated genes with antibodies against the P-TEFb subunits CycT and Cdk9 and the co-activator BRD4/fs(1)h. **b)** ChIP-qPCR validation of tissue-specific enrichment of CycT at DV regulated gene promoters (*dpp*, *tld*, *sog*, *ths*, *ind*, *sna* and *twi*) in *Toll* mutants. Enrichment is measured at DV targets relative to the signal at representative intergenic regions. Error bars show SEM. Significant differences in enrichment at promoters between the mutant that expresses the gene versus the mutants that do not (two tailed, unpaired t-test) are indicated by asterisks (* = *P* < 0.05, ** = *P* < 0.01, *** = *P* < 0.001). **c)** Metagene profiles and heatmaps (± 5 kb of CBP peak) of *Toll* mutant CycT, Cdk9 and BRD4/fs(1)h CUT&Tag signal (RPKM) at dorsal ectoderm, neuroectoderm and mesoderm enhancers. **d)** RT-qPCR quantification of *doc1, doc2, doc3* and *Elba3* mRNA levels (relative to *RpL32*) from doc enhancer (enh) del^Δ/Δ^ embryos (2-4 h AEL) and *PCNA-eGFP* and enh^+/+^ embryos (*n* = 3). **e)** ChIP-qPCR showing the enrichment of H3K27ac, H3K27me3 and CBP at the promoters of *doc1*, *doc2* and *doc3* in doc enh del^Δ/Δ^ embryos (2-4 h AEL) relative to enh^+/+^ embryos (*n* = 3-4). Relative occupancy is also shown at an intact doc enhancer (E4). Error bars show SEM. Significant differences in occupancy (two tailed, unpaired t-test) are indicated by asterisks (* = *P* < 0.05). **f)** ChIP-qPCR showing the % input precipitated by anti-CycT at the promoters and enhancers of DV regulated genes (*dpp*, *ind* and *twi*) and an intergenic control region in chromatin from wild-type and P-TEFb maternally overexpressed (OE) embryos (2-4 h AEL). Error bars show SEM. Significant differences in the % input precipitated at targets between wild-type and P-TEFb OE embryos (two tailed, unpaired t-test) are indicated by asterisks (* = *P* < 0.05). **g)** Odds ratios measuring the strength of association between *in situ* ectopic expression observed in P-TEFb OE, relative to wild-type embryos, for DV regulated genes. **h**) Correlation between *in situ* ectopic expression of DV genes in P-TEFb OE embryos (odds ratio P-TEFb OE/wild-type) and the relative CUT&Tag promoter occupancy of CycT at the same DV genes in inactive Toll mutant embryos relative to active. *P-*values are from Fisher’s exact test.

**Figure S5. Repressors occupy DV enhancers and promoters in a tissue-specific manner. a)** Overlap (%) of DV enhancers and promoters, partitioned by the tissue of target gene activity, with peaks called from ChIP-seq for Sna ^14 13^ and ChIP-nexus for Dl ^41^. **b)** Metagene profiles (± 5 kb of CBP peak) of *Toll* mutant (2-4 h AEL) Dl CUT&Tag signal (RPKM) at Sna repressed enhancers (*n* = 13). **c)** Metagene profiles (± 5 kb of CBP peak) of *Toll* mutant (3 h AEL) ATAC-seq signal (RPKM) at Sna repressed enhancers (*n* = 13). **d)** Metagene profiles (± 5 kb of CBP peak) of Dl ChIP-nexus and Sna and Pc (PRC1) ChIP-seq signal (RPKM) at DV enhancers and promoters. **e)** Metagene profiles comparing Pc ChIP-seq signal at DV enhancers and promoters partitioned by the tissue of target gene activity. **f)** Overlap (%) of DV enhancers and promoters with known Polycomb Response Elements (PREs) ^58^.

**Figure S6. Temporal dynamics of DV enhancer and promoter chromatin states. a)** Overlap (%) of DV and non-DV enhancers and promoters with Zld ChIP-seq peaks from nc 8, 13 and 14 wild-type embryos ^62^. For measuring overlaps, promoter regions were defined as (TSS ± 750 bp). **b)** (Top) Metagene plots and (bottom) heatmaps of Zelda (Zld) ChIP-seq enrichment in nc 8, 13 and 14 wild-type embryos at DV enhancers and promoters ^62^. **c)** Metagene plots of ChIP-seq enrichment for the pioneer factors Zld ^61^, opa ^63^, CLAMP ^64^ and GAF ^13,14^ from wild-type embryos at DV enhancers (± 5 kb of CBP peak) and promoters (± 5 kb of TSS). **d)** Images of whole-mount *in situ* hybridization with a probe against *schnurri* (*shn)* mRNA in wild-type and *Toll* mutant embryos (2-4 h AEL). **e)** Genome browser shot of ATAC-seq signal from wild-type naïve embryos (nc 11, 12 and 13) ^60^ and *Toll* mutant embryos (3 h, 4 h and 5 h AEL) alongside Zld, GAF, opa and CLAMP ChIP-seq and *Toll* mutant PRO-seq (3 h AEL) at the *shn* locus. The major *shn* promoter active during early embryogenesis and associated enhancers are denoted. **f)** (Top) Images of whole-mount *in situ* hybridization of LacZ reporter activity driven by VT enhancer fragments that overlap *shn* enhancers with predicted early (E1) and late (E2) embryonic activity, alongside *in situ* hybridization with a probe against endogenous shn mRNA in wild-type embryos at corresponding developmental stages. (Bottom) Plots of the mean ATAC-seq signal (log_2_ read count) at the E1 and E2 enhancers (3 h, 4 h and 5 h AEL) in the *gd^7^* mutant compared to *Toll^rm9/rm10^* and *Toll^10B^* (*n* = 3). Error bars show SEM. Significant differences in the accessibility between *gd^7^* and *Toll^rm9/rm10^*/*Toll^10B^* (two tailed, unpaired t-test) are indicated by asterisks (* = *P* < 0.05, ** = *P* < 0.01, *** = *P* < 0.001). **g)** Metagene plots of the ChIP-seq enrichment of histone marks (H3K27ac, H3K18ac, H4K8ac, H3K9ac, H3K4me1 and H3K4me3) ^66^ at DV, non-DV and shuffled enhancers and promoters from wild-type embryos at nc 8, 12 and 14 (early and late). **h)** Venn diagrams of the overlap between DV enhancers bound by H3K27ac, H3K18ac and H4K8ac across the time course. **i)** Overlap of DV enhancers acetylated or non-acetylated already at nc 8 with nc 8 Zld ChIP-seq peaks. **j)** Boxplots of CUT&Tag enrichment (log_2_ TPM) of CBP, Cdk9 and BRD4/fs(1)h at DV and non-DV enhancers and promoters relative to shuffled genomic control regions from wild-type embryos at nc 7-9, 11-13 and 14. P-values (Wilcoxon rank-sum test) show significant enrichment compared to shuffled regions. **k-l)** Metagene plots of **(k)** CBP-catalyzed histone marks from nc 8 ChIP-seq and **(l)** CBP CUT&Tag and ATAC-seq enrichment at the promoters of DV genes linked to early acetylated or non-acetylated enhancers at nc 8.

**Figure S7. Identification of DV relevant cell clusters from scRNA-seq data based on PRO-seq identified DV genes. a)** UMAP clustering of single-cell RNA-seq (scRNA-seq) data from wild-type embryos (2.5-3.5 h AEL) ^7^. DV-relevant clusters are shown in bold, **b)** Projections of the mean expression (z-score) of dorsal ectoderm, neuroectoderm and mesoderm genes identified by PRO-seq on the UMAP of wild-type cells from DV relevant clusters from **a** reclustered according to the expression of PRO-seq DV genes (see methods). **c)** Projections of the expression of the marker DV used to identify the 6 clusters from the UMAP from wild-type cells in **b** and violin plots of the expression (log_2_ TPM) for each gene in the assigned cell clusters (see Fig. 6a). **d)** The assignment (%) of DV relevant cells from wild-type and *Toll* mutant embryo scRNA-seq UMAP clusters (see Fig. 6a). **e)** Projections of *dpp*, *ind* and *twi* expression on UMAPs from *Toll* mutant scRNA-seq (see Fig. 6a).

**Figure S8. Transcriptome-wide inference of burst kinetics from single-cell expression data. a)** Correlation plots of burst size (log_2_) and burst frequency (log_2_) kinetics inferred for all genes in DV relevant (dorsal ectoderm, neuroectoderm (early) and mesoderm (early)) scRNA-seq clusters between *PCNA-eGFP* and *w^1118^* control (wild-type) lines. *R*^2^ and *P*-values relating to the correlations are shown. **b)** Boxplots of the burst size (log_2_) and burst frequency (log_2_) for DV and AP regulated genes, alongside genes partitioned by the stage of expression during early embryogenesis. **c)** Correlation plots of burst kinetics and the PRO-seq promoter read count (log_2_) for DV regulated genes. **d)** Venn diagram showing the overlap between differentially expressed DV genes identified from *Toll* mutant PRO-seq (*n* = 195), the DV genes paired with enhancers (*n* = 105) and DV genes with a significant change in either one or both inferred transcriptional kinetic between DV relevant clusters from the wild-type embryo scRNA-seq data (*n* = 47). **e)** The number of DV genes with a significant kinetic change between DV-relevant clusters and the proportion (%) that change in burst size and frequency for the genes partitioned according to the tissue of expression. **f)** Heatmaps showing the expression level (z-score) of DV genes with a significant change in at least one kinetic parameter across profiled in single cells assigned to DV-relevant clusters, partitioned by the tissue of expression. Whether each DV gene changes significantly in burst size (BS) and/or burst frequency (BF) and has been paired to one or more enhancers is denoted. For each gene, the mean BF, BS, *k*_on_, *k*_off_ and *k*_syn_ kinetic values from DV-relevant clusters are plotted. **g)** Correlation plots of the fold change in burst kinetics (log_2_ active tissue/inactive tissues) and DV enhancer and promoter tissue-specific chromatin state scores. **h)** Boxplots showing the inferred transcriptional parameters for DV genes in the active tissue and inactive tissues. DV genes are partitioned into kinetic classes based on whether they have significant changes in burst frequency, burst size or both between the active and inactive tissues (see Fig. S8h). *P*-values (Wilcoxon rank-sum test) show significant differences in each parameter for the kinetic classes between the active and inactive tissues. **i)** Boxplots of the log_2_ fold change (active tissue/inactive tissues) in each parameter for the kinetic classes. *P*-values (Wilcoxon rank-sum test) show significant differences the fold change between different kinetic classes. **j)** Heatmap (from 0 to 1) showing the coefficient of determination (*R^2^*) for DV enhancer and promoter tissue-specific chromatin state compared to kinetic parameter scores. DV genes were classified according to whether they showed a significant change in BS, BF or both between their tissue of activity and inactive tissues. Comparisons with significant positive and negative correlations are denoted by boxes.

