## Supplementary figures and images for "Comprehensive interrogation of a *Drosophila* embryonic patterning network reveals the impact of chromatin state on tissue-specific burst kinetics and RNA Polymerase II promoter-proximal pause release"

### Supplemental Figures

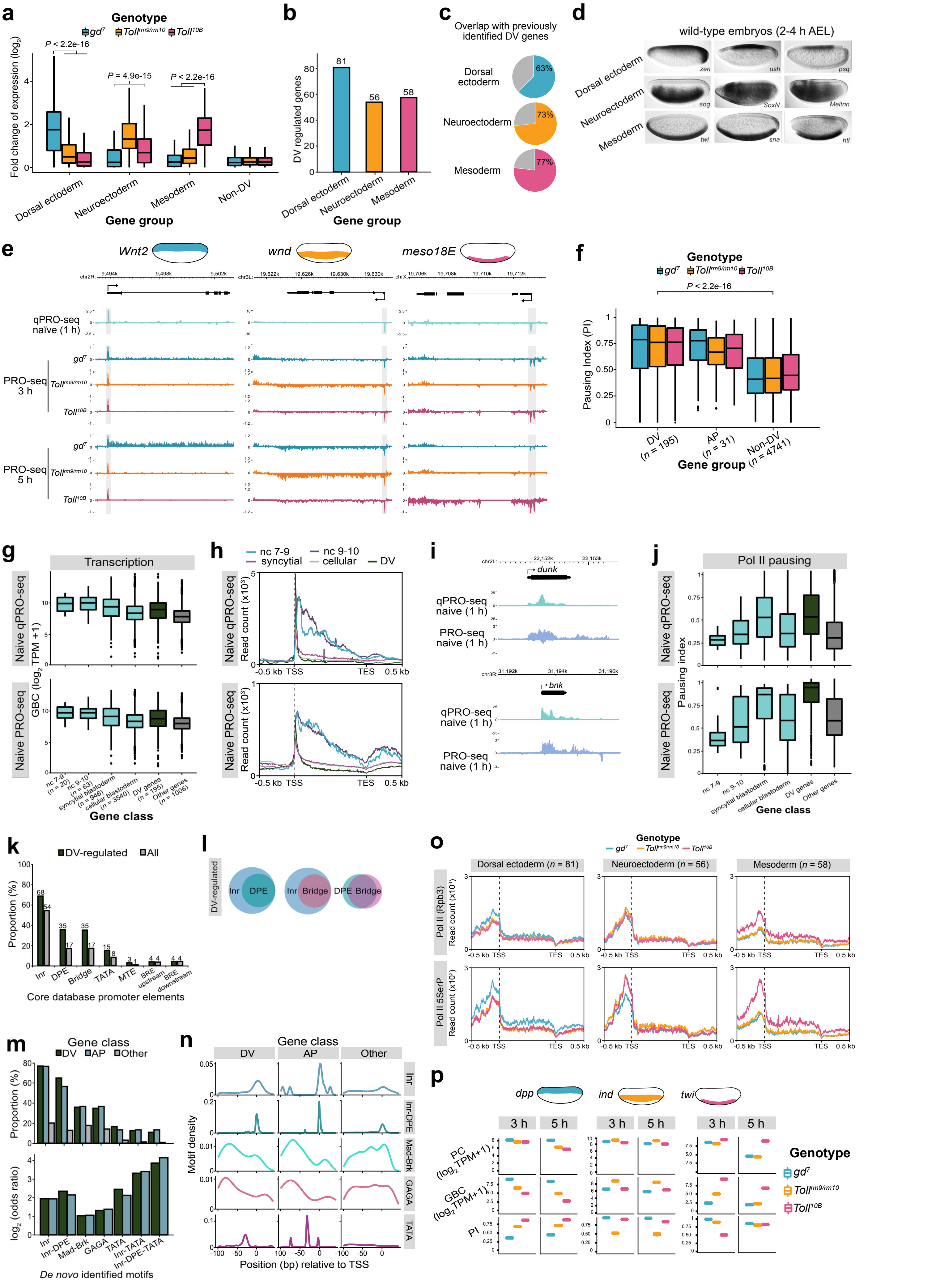

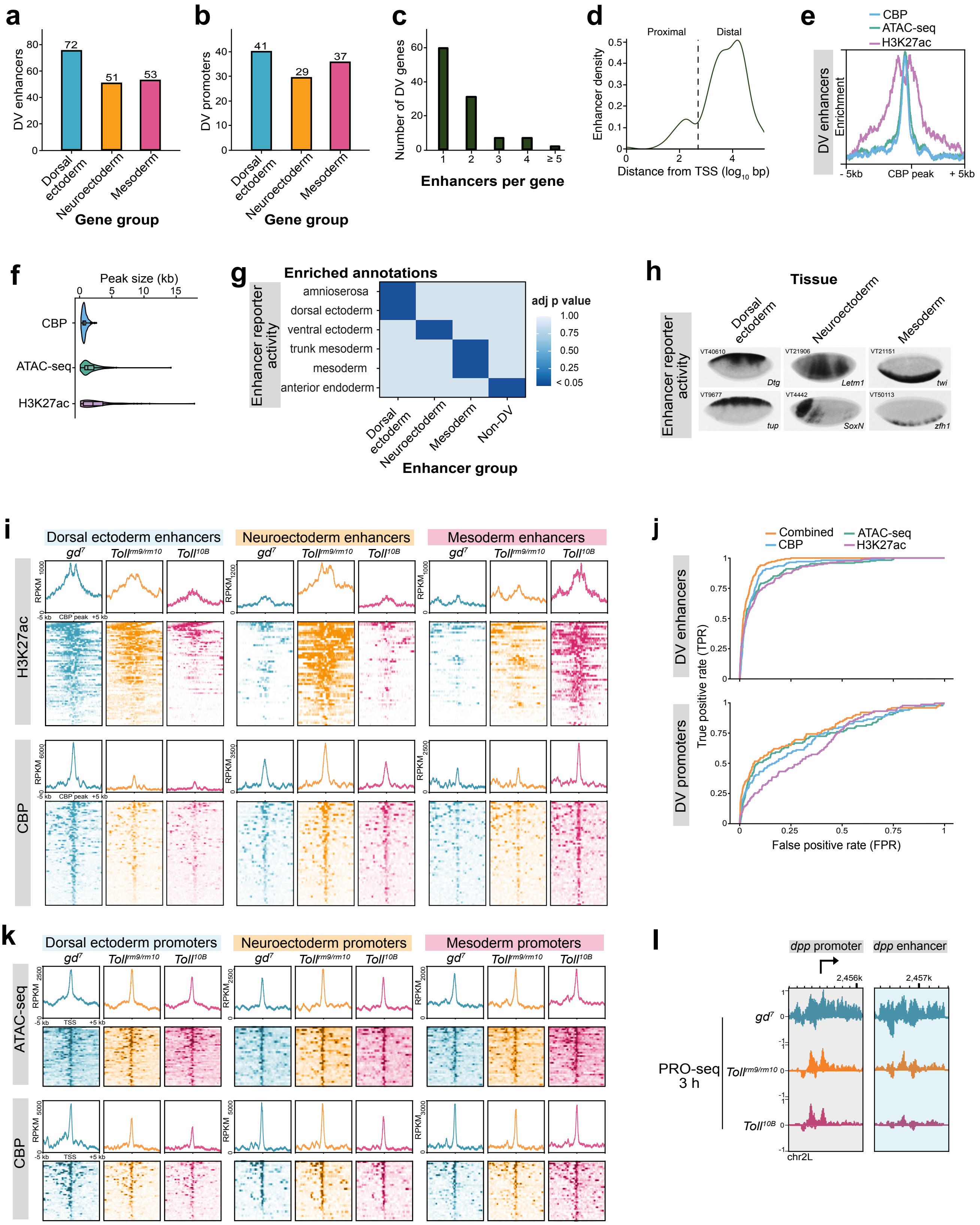

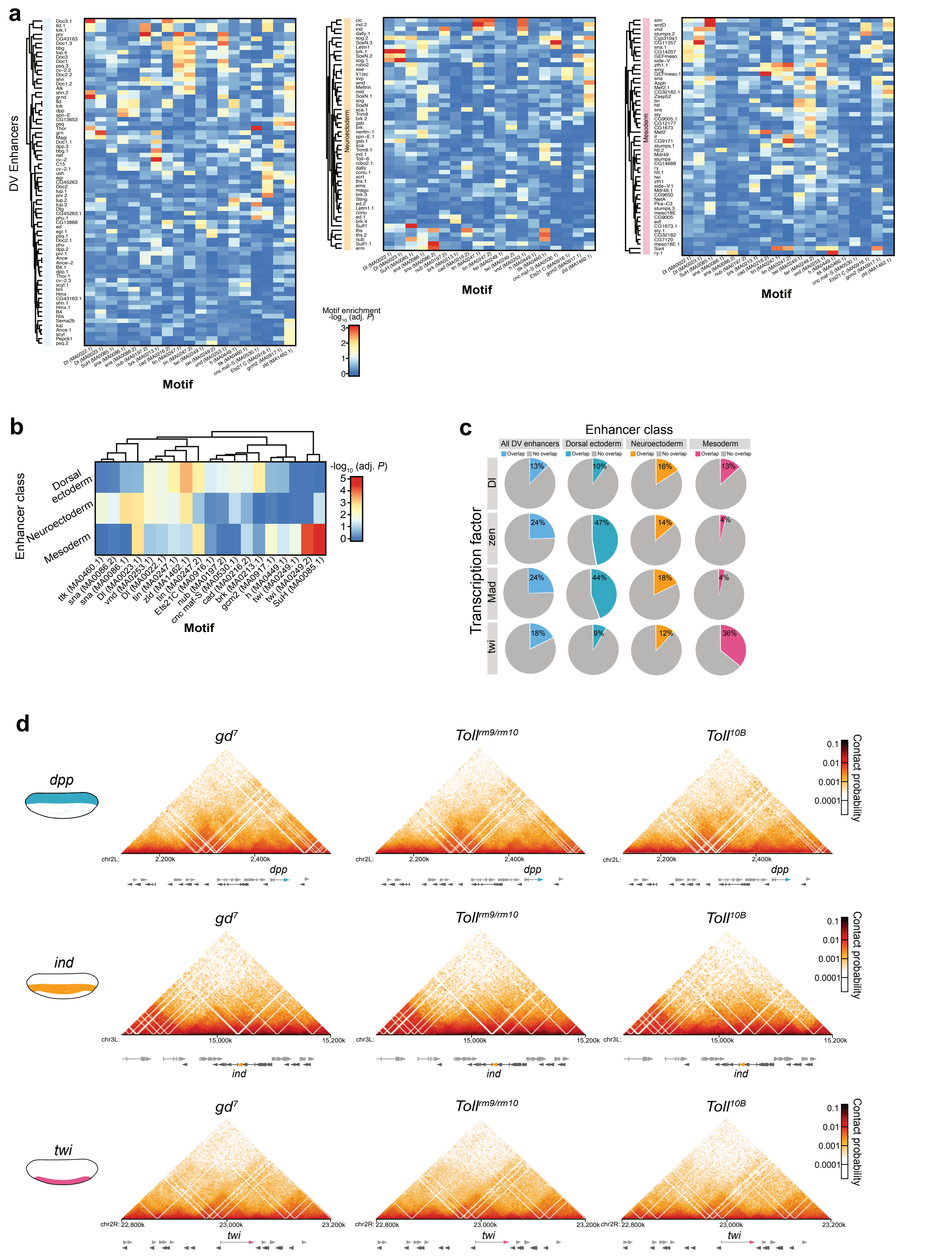

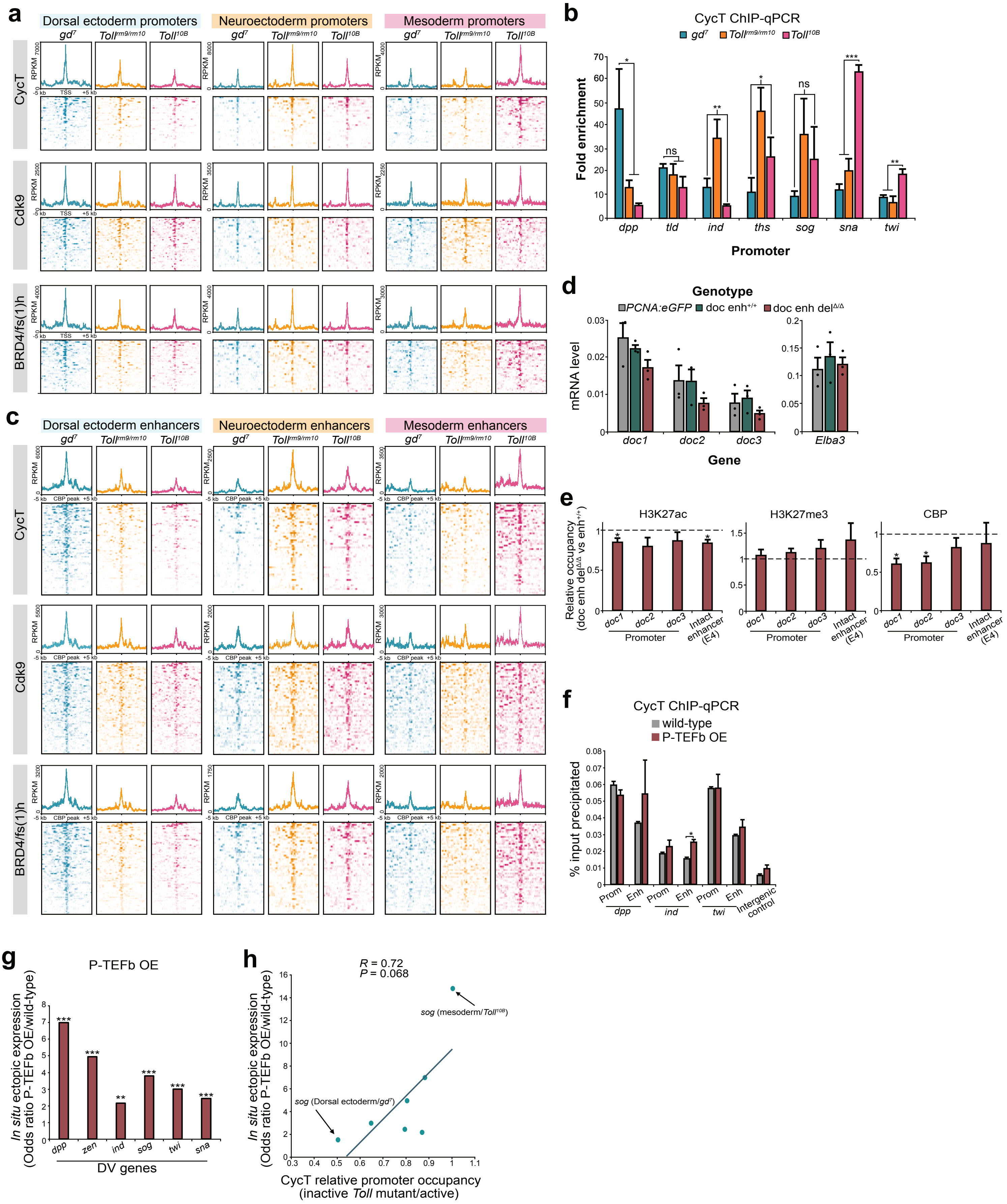

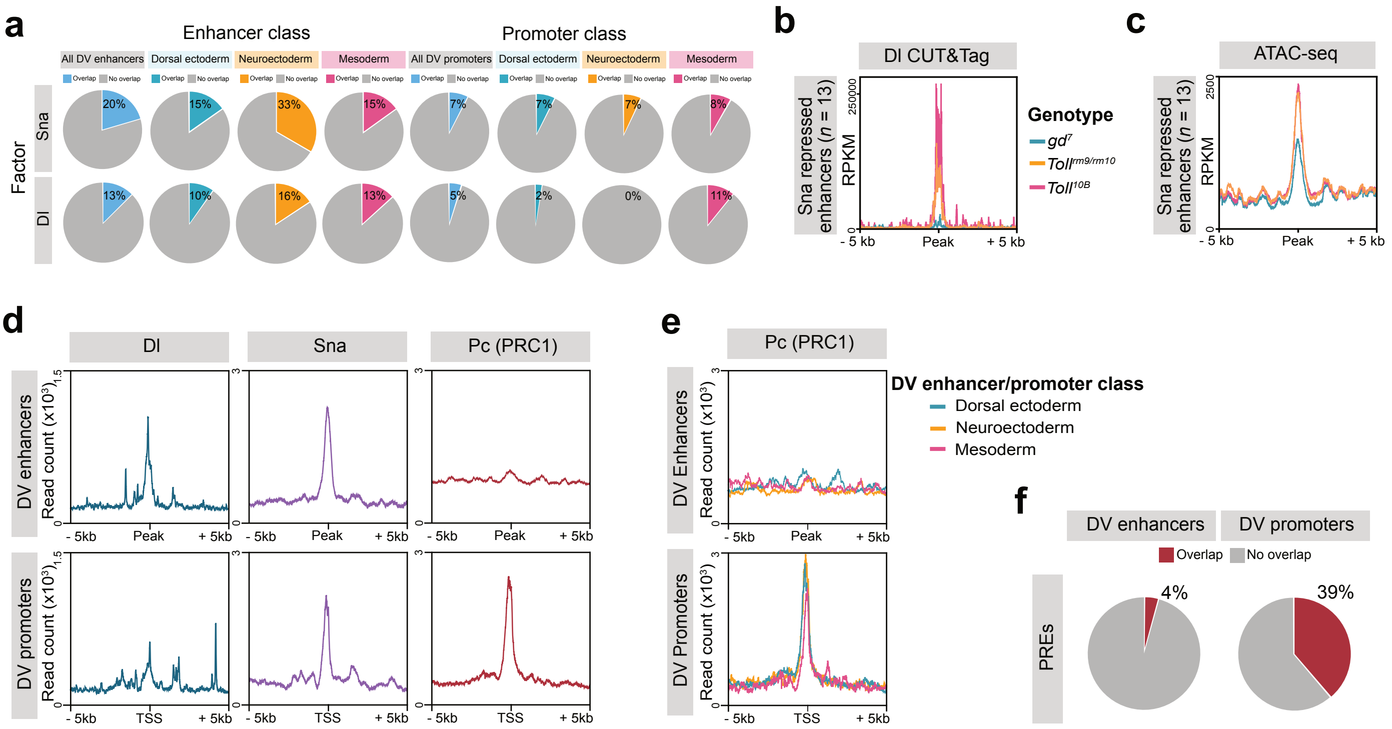

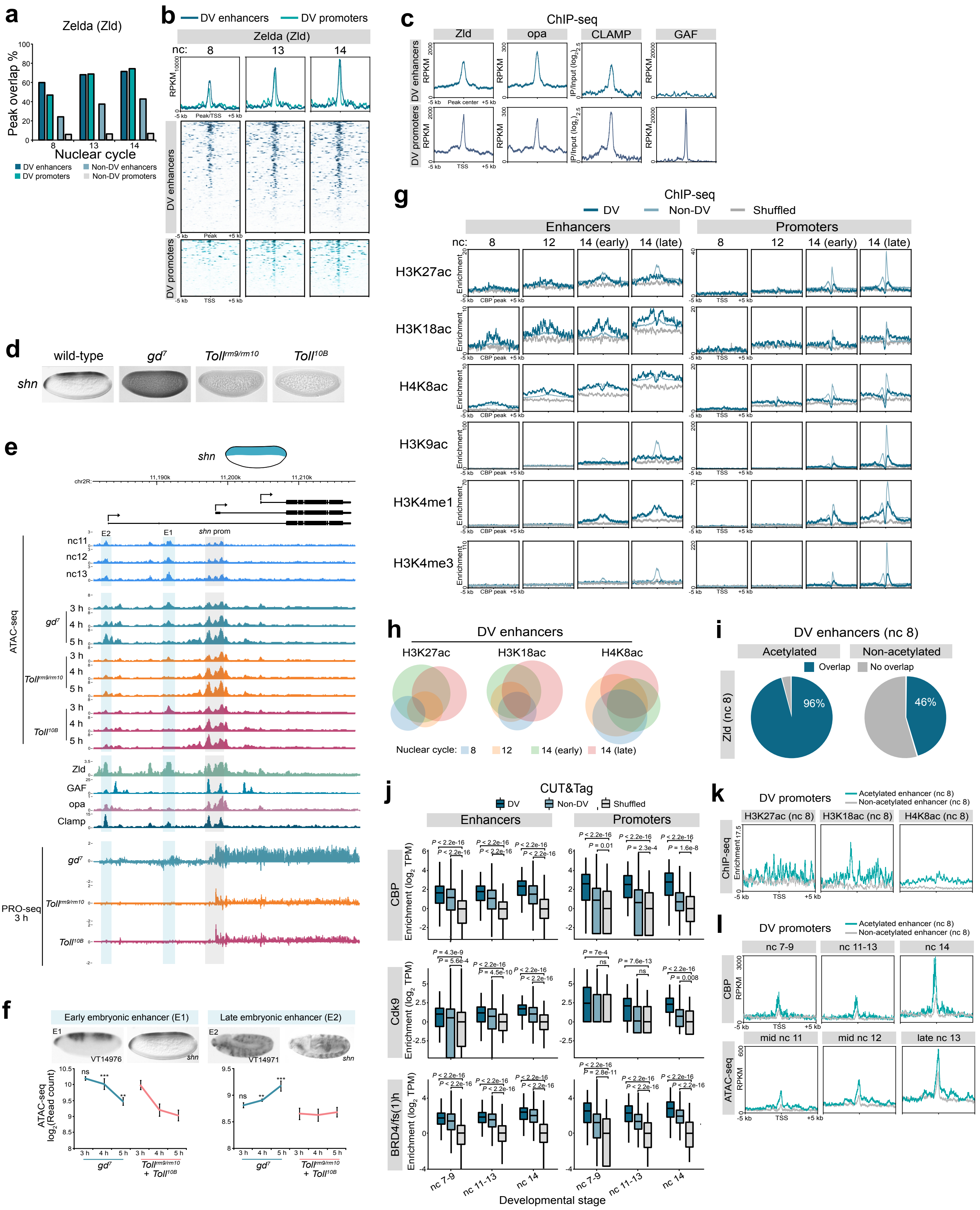

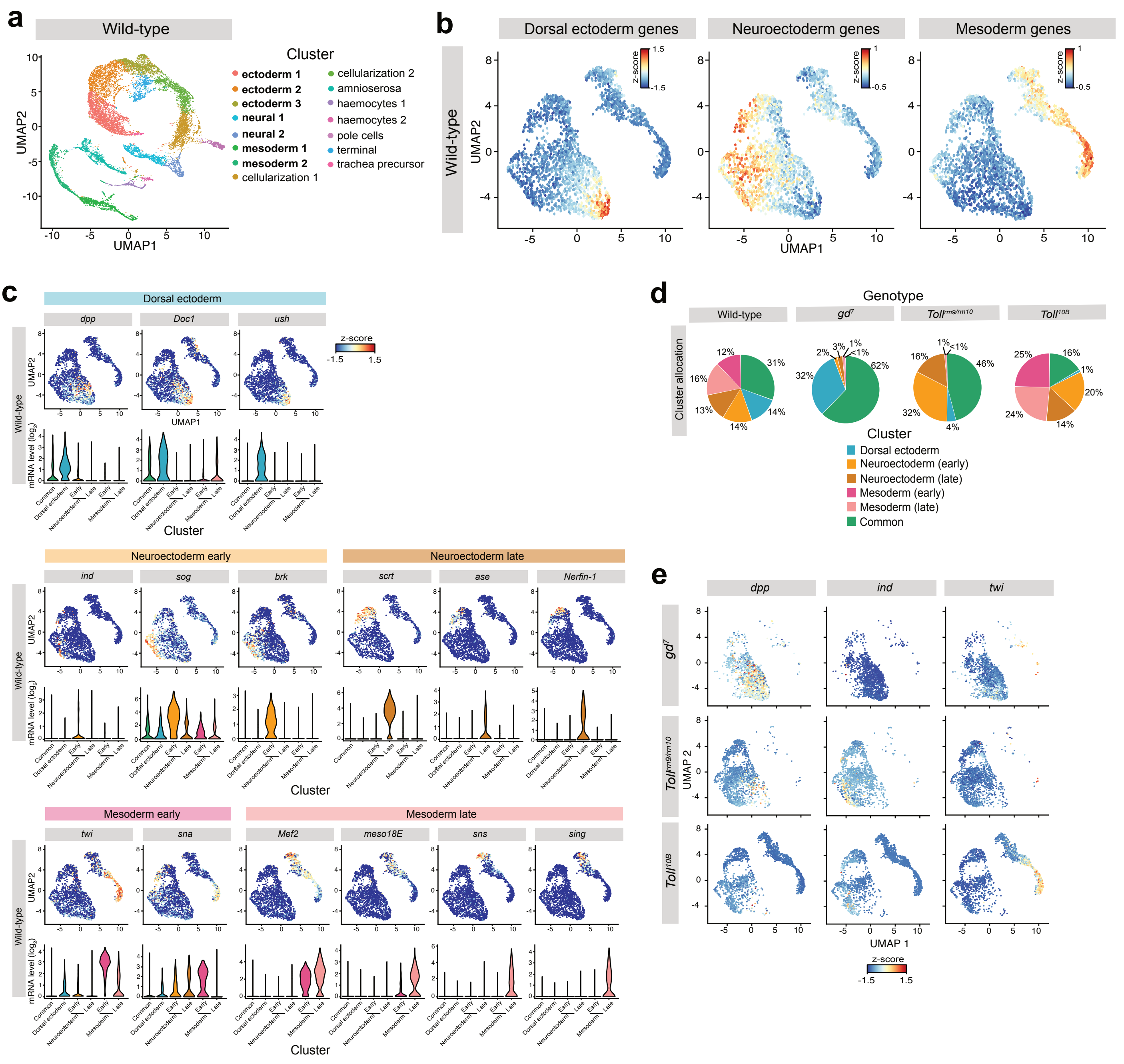

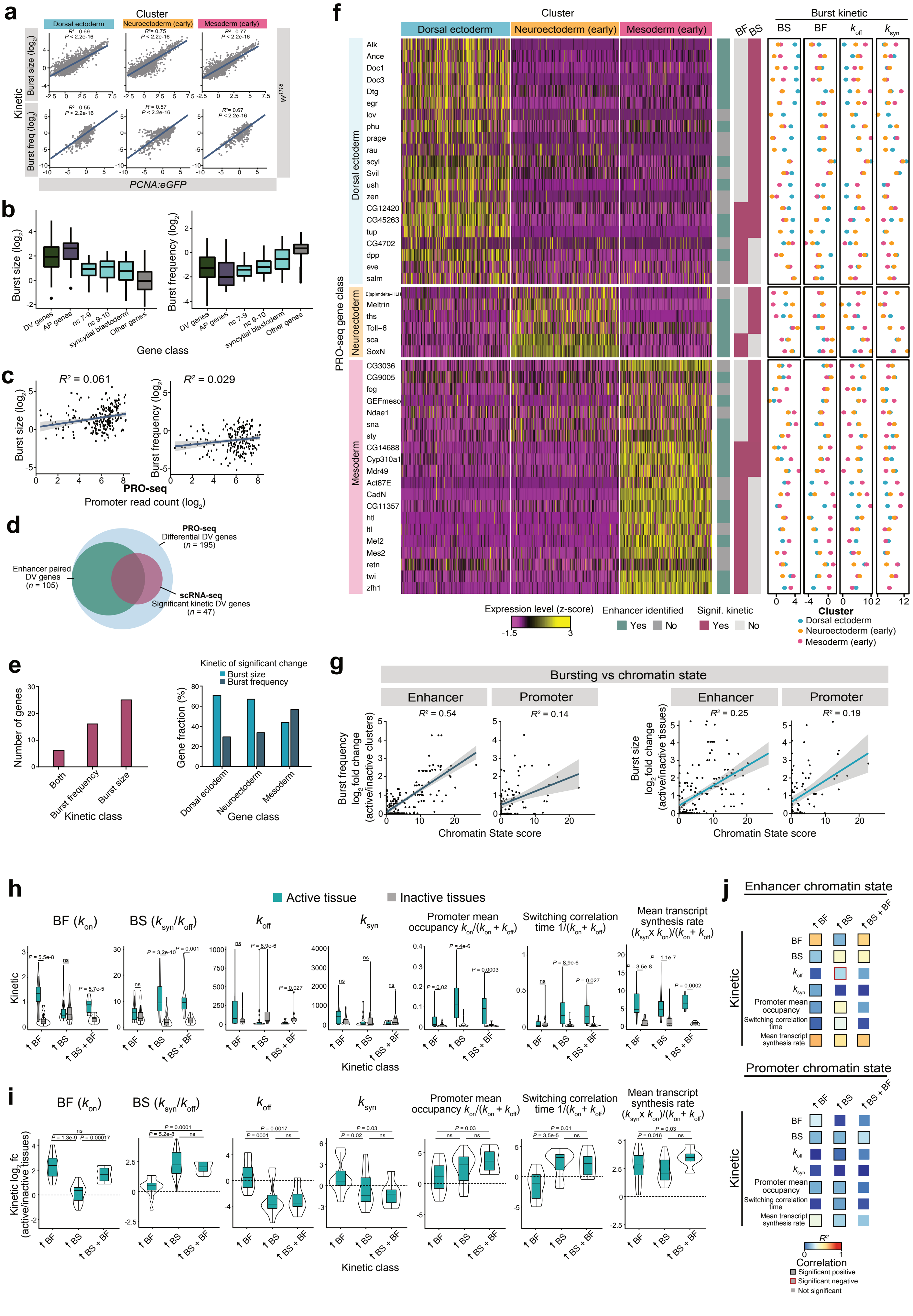
